## Supplementary Figures with legends for "Identification of a Rabenosyn-5 like protein and Rab5b in host cell cytosol uptake reveals conservation of endosomal transport in malaria parasites"

A

**FYVE domains**

**Logo of motifs**

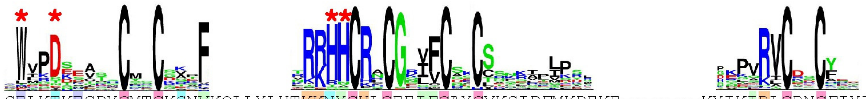

```
Pf_Rbsn5L          44 SELKTKEGDYCMTCKSNVKQLLYLHTKKNYCHLCEEIFCAYCVKSIDFMKDEKE-------KYIKIRLCRNCFIYIN  119
PF3D7_1460100      32 LWVPDEEVTNCYSCNVVF----NVRVRKHHCRACGNVFCSNCSDNKIKISEYSYS--------EKVRVCDRCFMERS   96
human_Protrudin   122 KRYPTNNFGNCTGCSATF----SVLKKRRSCSNCGNSFCSRCCSFKVPKSSMGAT--APEAQRETVFVCASCNQTLS  192
yeast_PEP7        206 PWRDDRSVLFCNICSEPF----GLLLRKHHCRLCGMVVCDDANRNCSNEISIGYL-20aa-IPISIRLCSHCIDMLF  297
human_EEA1       1298 KWAEDNEVQNCMACGKGF----SVTVRRHHCRQCGNIFCAECSAKNALTPSSKK----------PVRVCDACFNDLQ 1411
human_Rbsn5       147 PWVNDQDVPFCPDCGNKF----SIRNRRHHCRLCGSIMCKKCMELISLPLANKLT-43aa-DDDRIRCCTHCKDTLL  260
human_FYV1        148 YWMPDSQCKECYDCSEKF----TTFRRRHHCRLCGQIFCSRCCNQEIPGKFMGYT--------GDLRACTYCRKIAL  218
```

Positively charged
Negatively charged
Polar Uncharged
Hydrophobic
Cysteine
* residues critical for PI3P binding that do not follow consensus in Pf_Rbsn5L

B

**Rab-binding domains (RabBD)**

```
                                  +      +               ++      +
PF_Rbsn5           130 ----SVDRKAIDLQNSFNDISNCYTNLCSNVPQLNGLVLLCENNKEFLDSFKSEIKQLEE 195
yeast_PEP7         464 --------VLKE---QSYLI-ESTIQDYKKQRRLEEIVTLNKNLEELHSRIH-------- 503
Drosophila_Rbsn5   441 ASKSSLDDPLIE---QINII-KGYIKQARQDMNFEVVETLELNLRELQREVYERQRQS-- 494
human_Rbsn5-copy1  458 --------PLLQ---QIHNI-TSFIRQAKAAGRMDEVRTLQENLRQLQDEYDQQ------ 499
human_Rbsn5-copy2  739 ---------LLQ---QIDNI-KAYIFDAKQCGRLDEVEVLTENLRELKHTLAKQ------ 779
                                  :   .   *            .:: :  *  * .::
```

Positively charged
Negatively charged
Polar Uncharged
Hydrophobic
Cysteine
+ involved in conserved hydrogen bonds in human Rbsn5

C

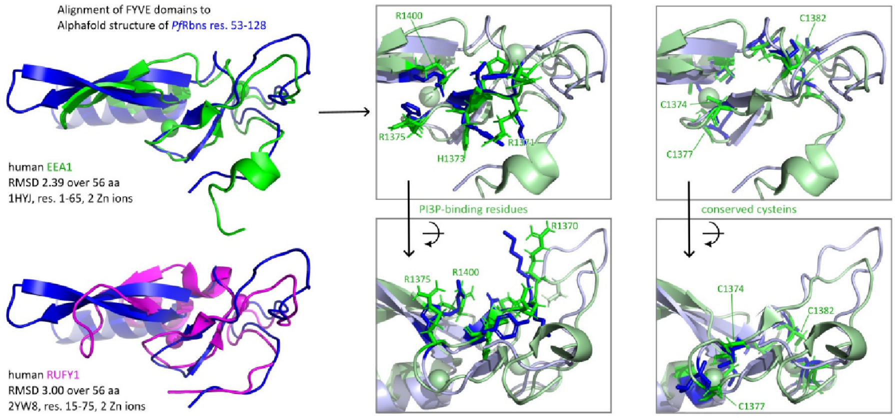


D

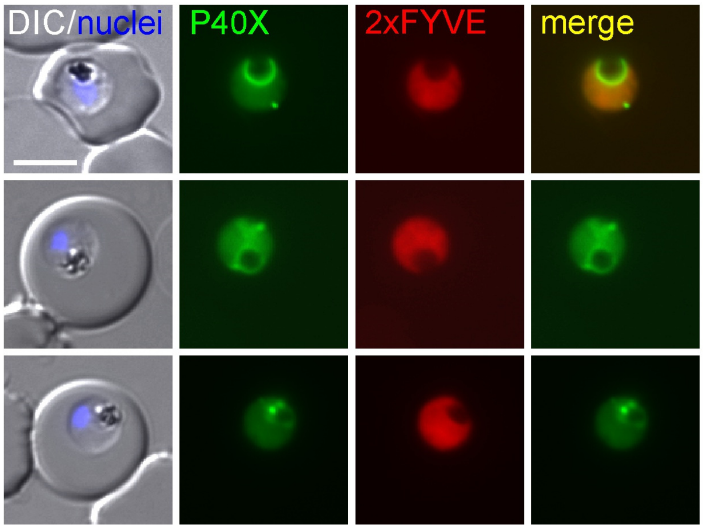


**Figure S1. Domain motifs in PfRbsn5. A-B,** Multiple sequence alignment of (A) FYVE and (B) Rab-binding domains of the indicated proteins and regions. Residues indicated as conserved were colored according to their physical properties. Sequence logo of the conserved residues in A was generated from the sequences of all FYVE domains found in *H. sapiens, S. cerevisiae, T. brucei, A. thaliana, G. intestinalis,*

*P. falciparum* and *T. gondii*. Red asterisks show amino acids important for PI3P binding or specificity that are not conserved in the FYVE domains of PfRbsn5L and human Protrudin (which contains a FYVE domain not binding PI3P but other phosphoinosities) while they are conserved in PF3D7_1460100 (FCP), the likely PfEEA1 [36]. In B, similarity according to clustal omega (dots and asterisks) are indicated below the alignment; residues involved in hydrogen bonds that are conserved between both Rab-binding domains of human Rbsn5, according to PDB 1z0k and 1z0j, were marked with + above. **C**, Structural alignment of AlphaFold2 predicted structure of PfRbsn5L FYVE domain compared to the experimental structures of human EEA1 and human RUFY1. **D**, Expression of a tandem of the PfRbsn5L FYVE (2xFYVE) domain fused to mCherry in parasites expressing P40X-GFP (P40X) to mark PI3P positive regions. Nuclei were stained with DAPI; DIC, differential interference contrast; merge: overlay of green and red channel; size bar: 5μm.

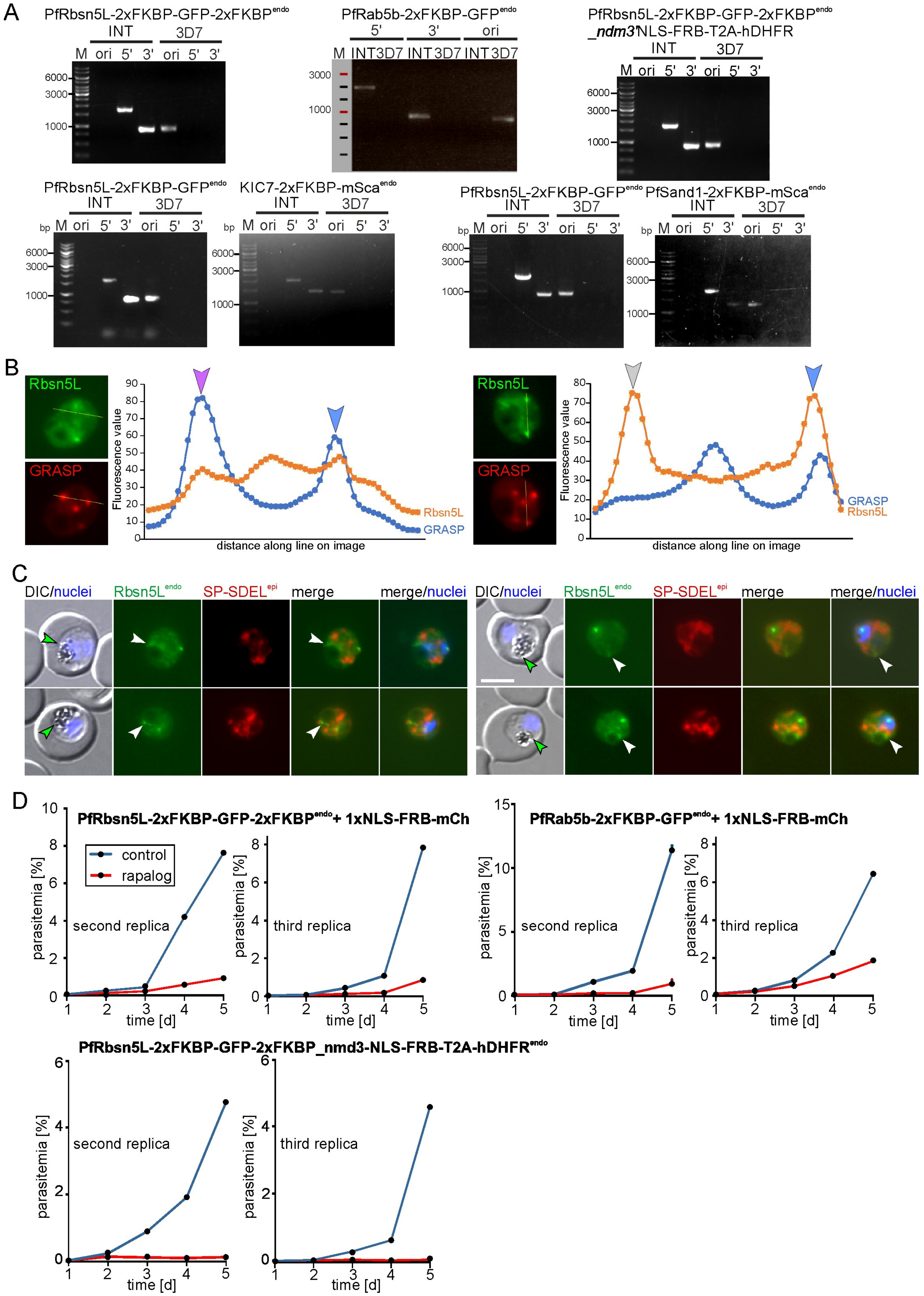


**Figure S2. Confirmation of correct genomic integration and additional flow cytometry growth curves. A,** Agarose gels showing PCR-products to assess correct integration of the SLI-plasmids into the genome of *P. falciparum* (3D7) parasites to obtain the indicated cell lines**.** 3D7, parent cell line; INT, integration cell line; 5', PCR product across the 5' integration junction; 3', PCR product across the 3' integration junction; ori, original locus (absence showing lack of parasites with unmodified locus). M, marker with selected fragments indicated in bp. Primers used for integration confirmation and expected sizes of PCR products listed in Table S1 and sequences of the plasmids in File S1. **B,** Intensity profiles along the indicated lines in images from Fig. 1C generated with ImageJ. Lines were drawn in both images using the synchronize images command and the plot profile values were used to draw graphs in Excel. Arrows show green foci and are color coded as in Fig. 1C. **C**, Live-cell microscopy images of PfRbsn5L-2xFKBP-GFP-2xFKBP$^{endo}$ parasites, co-expressing the ER-marker STEVOR-SP-mScarlet-SDL$^{epi}$ (SP-SDEL$^{epi}$). White arrows show PfRbsn5$^{endo}$ foci close to the food vacuole that were added at the same position in the DIC/Hoechst image as green arrows to illustrate the position relative to the food vacuole. **D,** Replicates of the flow cytometry growth curves shown in the main figures (cell lines indicated).

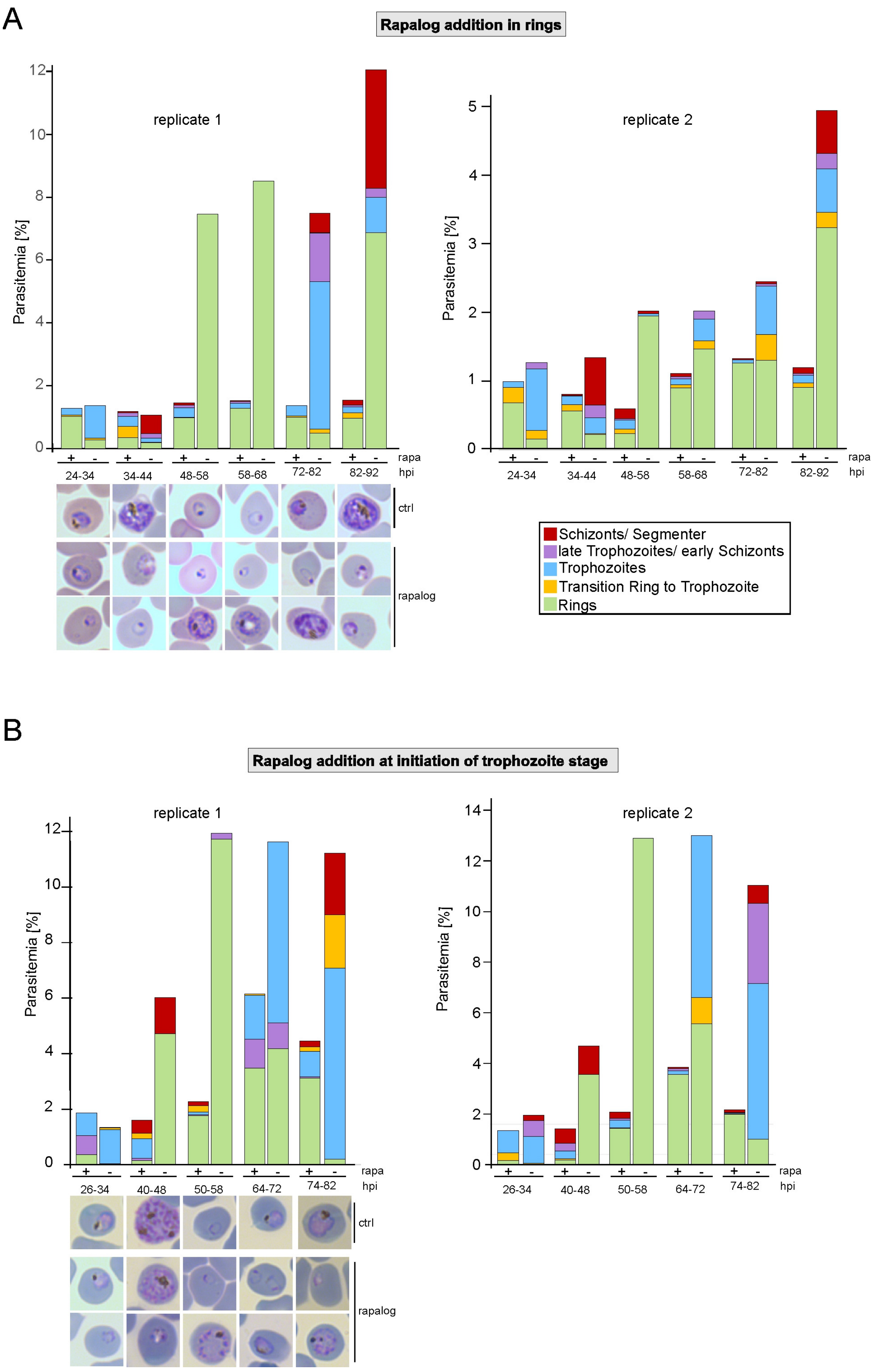


**Figure S3. Inactivation of PfRbsn5L in synchronous parasites. A-B,** Synchronous parasites were inactivated in rings (A) or at the start of the trophozoite stage (B) (see materials and methods) and the development monitored based on Giemsa smears. The rings monitored correspond to parasites where PfRbsn5L was inactivated 0 – 8 h post invasion (hpi), the trophozoites at 16 – 24 hpi (n = 2 independent experiments; Giemsa smears shown only for replicate 1). Note that cells scored as rings after prolonged growth arrest in + rapa in the ring induction showed various morphological alterations not specifically scored.

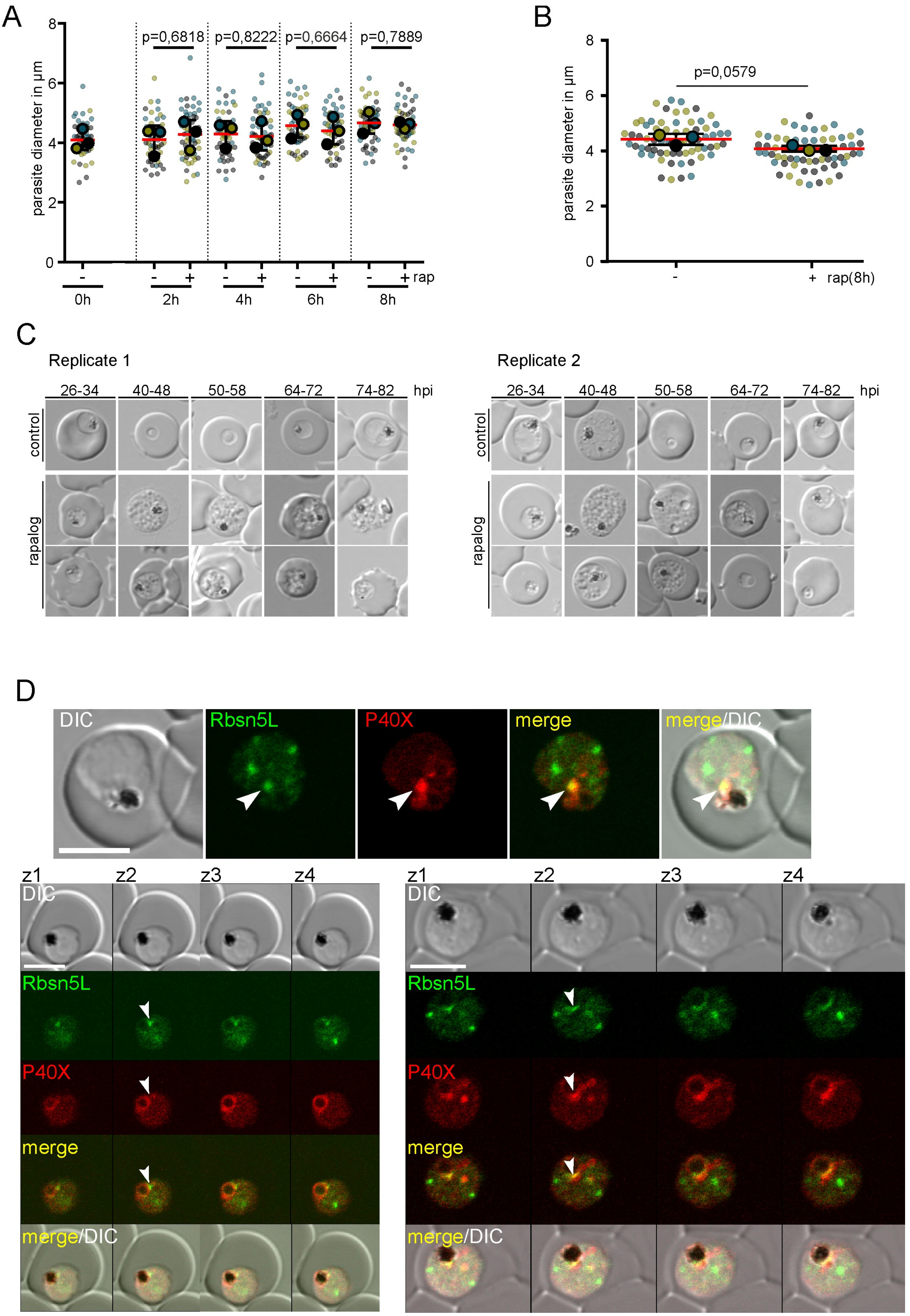


**Figure S4. Additional data for bloated food vacuole and vesicle accumulation assays and vesicle accumulation phenotype and co-localization of PfRbsn5L with regions containing PI3-P. A-B,** Superplots showing diameter in µm of parasites analyzed in the vesicle accumulation assay in Figure 1G

(A) and of parasites analyzed in the bloated FV assay in Figure 2B (B). Parasites from n = 3 independent experiments are distinguished by blue, yellow, and black dots; two-tailed unpaired t-test of the means, p-values indicated; mean (red bar); error bars (black) show S.D.. **C**, DIC example images showing the phenotype of time points of the parasites from the stage growth assay in Figure S3 that resulted in the growth phenotype (time points indicated). **D**, Confocal microscopy images of PfRbsn5L$^{endo}$ parasites (PfRbsn5L: green channel) co-expressing the mScarlet tagged PI3P marker P40X (red channel) used for the experiments shown in Figure 2E. Top, single z-slice, bottom panels show 4 consecutive confocal z-slices (z1-4). White arrows: PI3P positive regions overlapping with PfRbsn5L accumulations adjacent or at the food vacuole. DIC, differential interference contrast; merge, overlay of red and green channel. Size bars, 5 μm.

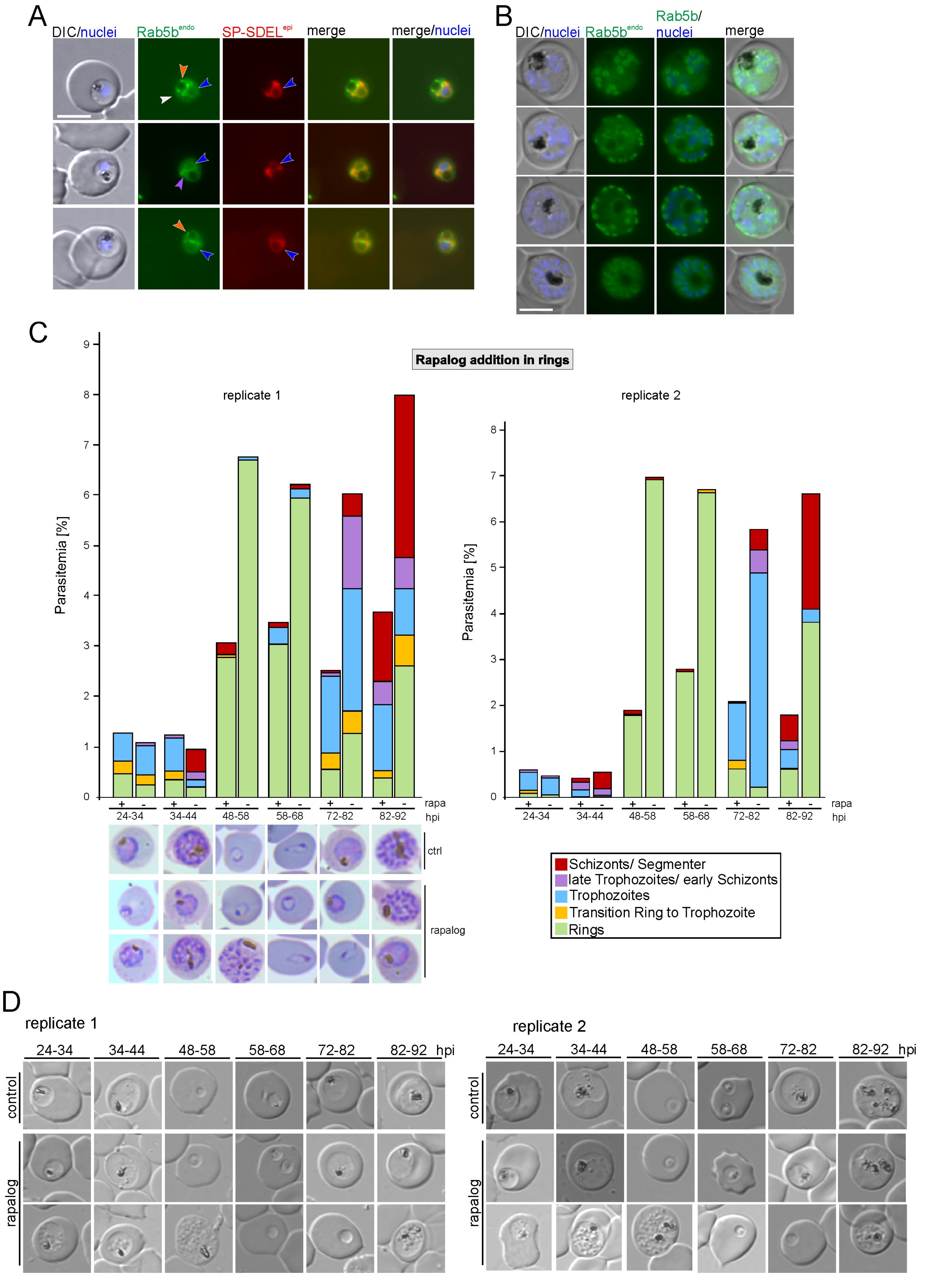


**Figure S5. Additional Rab5b localization data and stage-specific growth phenotype.**
**A, B,** Live-cell microscopy images of PfRab5b$^{endo}$ parasites co-expressing the ER marker STEVOR-SP-mScarlet-SDL$^{epi}$ (SP-SDEL$^{epi}$) (A), and of PfRab5b$^{endo}$ schizonts (B) showing a location suggestive of the IMC. Merge, overlay of green and red channels. Arrows in A show: blue, ER membrane; white, parasite

plasma membrane; orange, Rab5b foci at the FV not overlapping with the ER marker; purple, possible FV loop-like structure. Size bar 5µm. Nuclei were stained with DAPI. **C**, Growth assay after conditional inactivation of Rab5b in synchronous rings monitoring progression through the cycle based on Giemsa smears (the parasites monitored correspond to parasites where Rab5b was inactivated 0 – 8 h post invasion (hpi)) (n = 2 independent experiments). Example Giemsa smear images shown for replicate 1. **D**, DIC example images showing the phenotype in the parasites from the stage growth assay in C of the time points (indicated) leading to the growth phenotype.

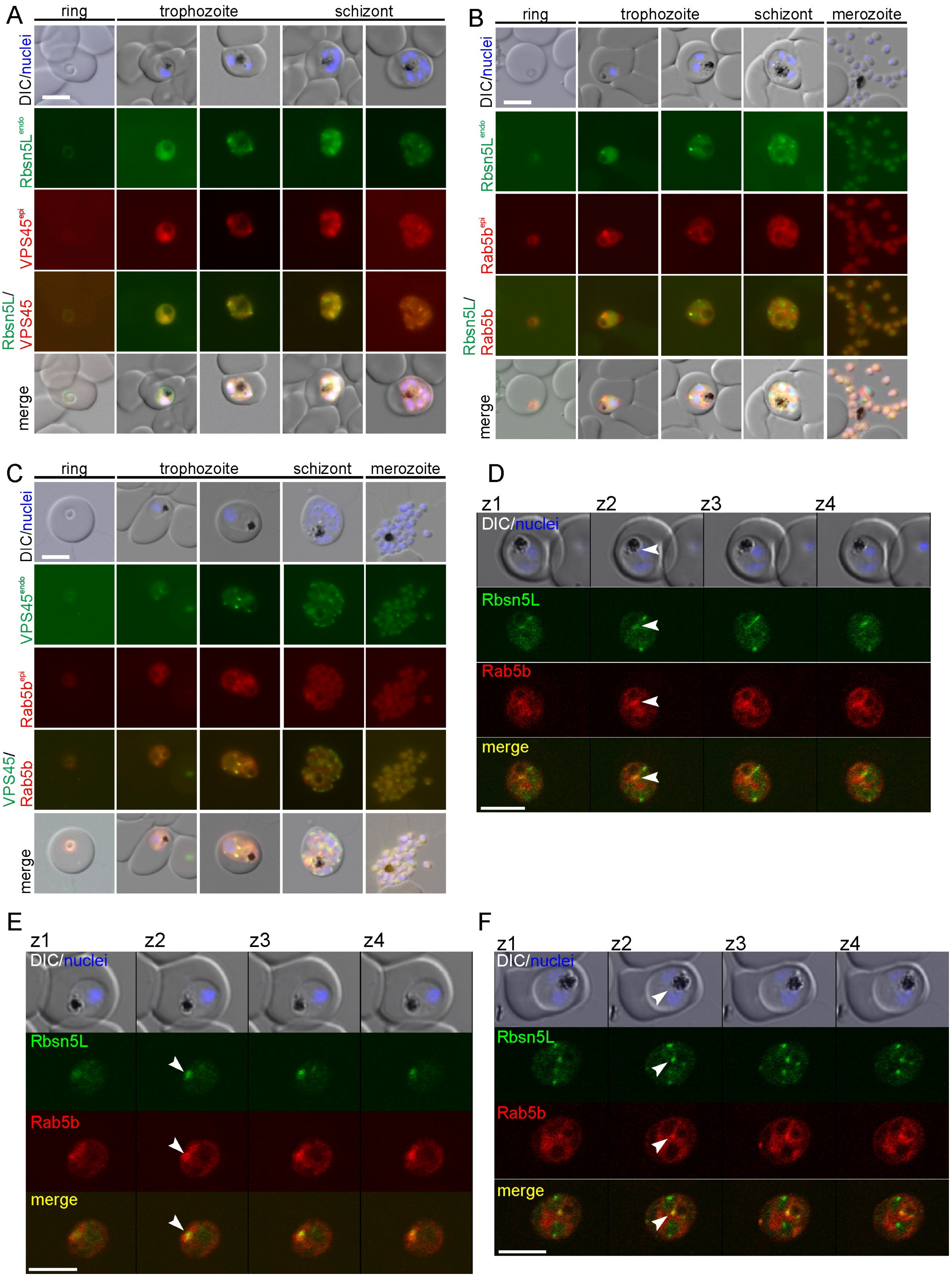


**Figure S6. PfRbsn5 and PfVPS45 co-localize with PfRab5b positive membranes. A-C,** Live-cell microscopy images of the indicated stages of PfRbsn5-2xFKBP-GFP-2xFKBP$^{endo}$ parasites, co-expressing

PfVPS45-mCh$^{epi}$ (A), PfRbsn5-2xFKBP-GFP-2xFKBP$^{endo}$ parasites, co-expressing PfRab5b-mCh$^{epi}$ (B), and PfVPS45-2xFKBP-GFP$^{endo}$ parasites, co-expressing PfRab5b-mCh$^{epi}$ (C). DIC, differential interference contrast; endo, endogenous; epi, episomal. Scale bar, 5 µm. Nuclei were stained with DAPI. **D-F**, Confocal microscopy images of PfRbsn5L$^{endo}$ + Rab5b$^{epi}$ parasites (PfRbsn5L: green channel; Rab5b: red channel) showing 4 consecutive confocal z-slices (z1-4) per cell. White arrows: Regions where PfRbsn5L overlaps with Rab5b. Nuclei were stained with Hoechst; DIC, differential interference contrast; merge, overlay of red and green channel. Size bars, 5 µm.

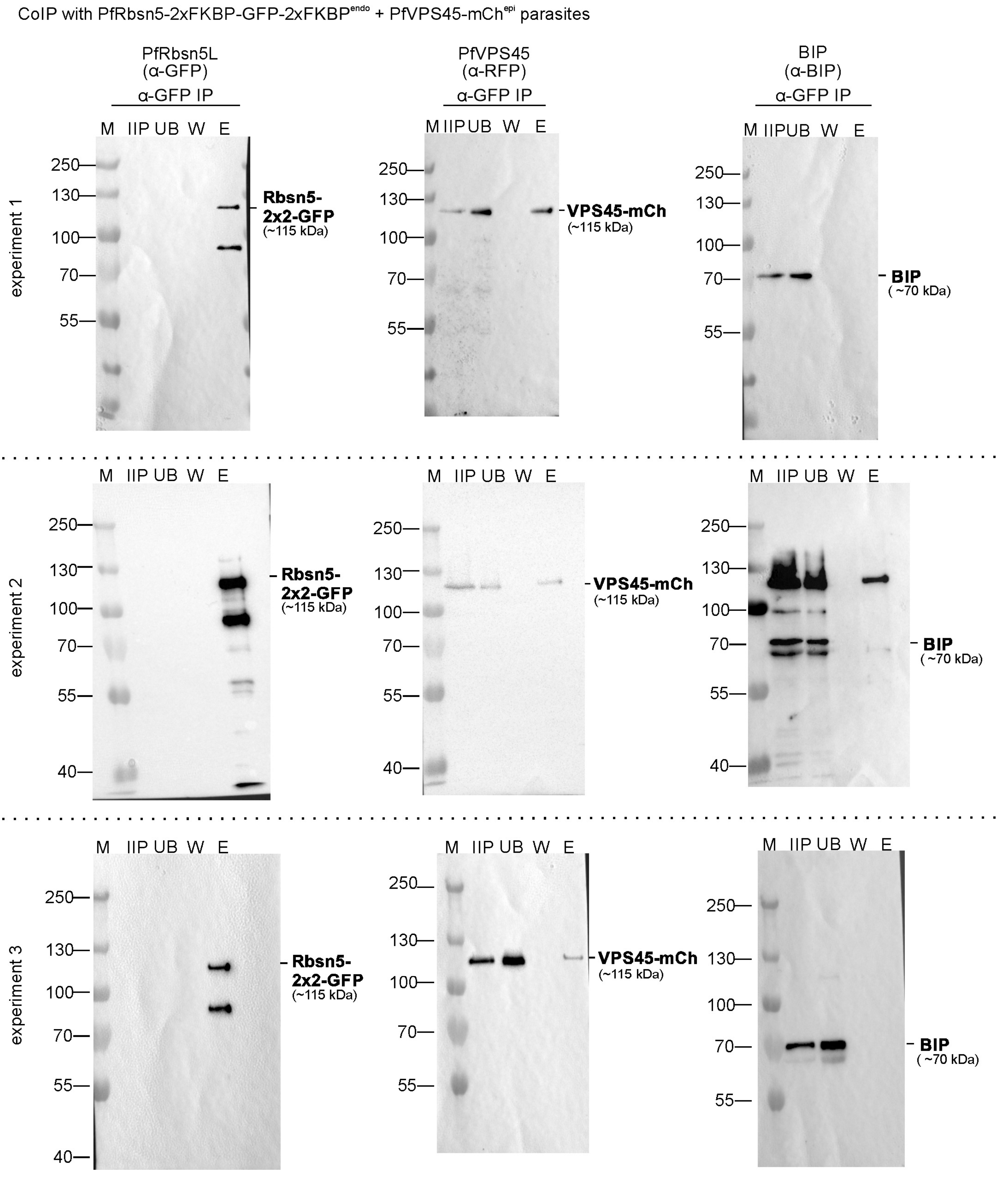


**Figure S7. PfVPS45 interacts with PfRbsn5.** Replicas and complete blots of 3 independent experiments of Immunoprecipitation (IP) of extracts from PfRbsn5$^{endo}$ parasites co-expressing PfVPS45$^{epi}$ (one cropped version is shown in Figure 5D). Note that only the indicated band on the anti-BIP blot in experiment 2 corresponds to BIP, the other bands are from a previous probing of the same membrane. IIP, IP-input extract; UB, unbound (total extract after IP); W, last wash; E, eluate; endo, endogenous; epi, episomal.

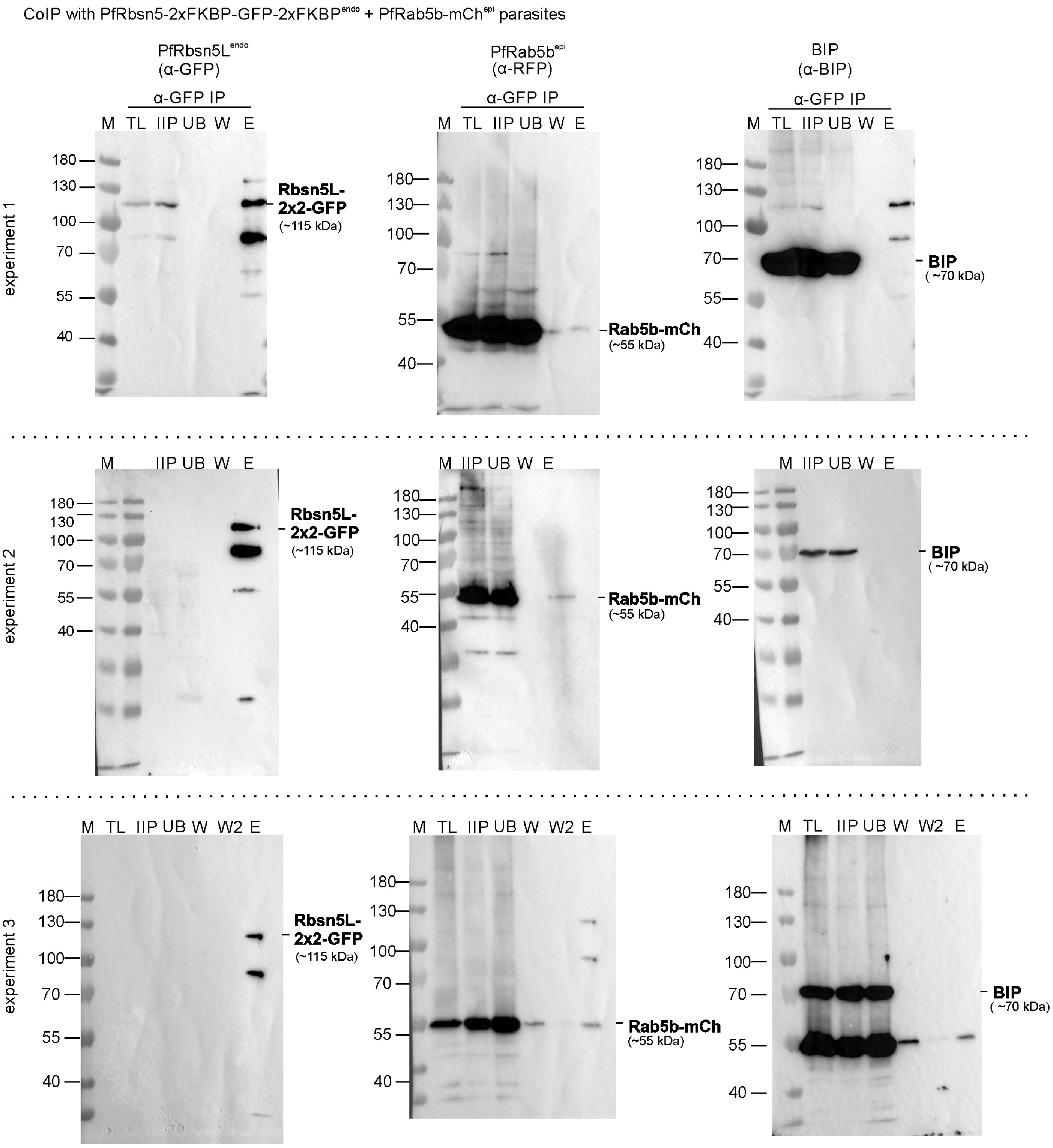


**Figure S8. PfRab5b interacts with PfRbsn5.** Replicas and complete blots of 3 independent experiments of Immunoprecipitation (IP) of PfRbsn5$^{endo}$ parasites co-expressing PfRab5b$^{epi}$ (one cropped version is shown in Figure 5E). Note that only the indicated band on the anti-mCherry and anti-BIP blots in experiment 3 correspond to the respective proteins, the other bands are from a previous probing of the same membrane. IIP, IP-input extract; UB, unbound (total extract after IP); W, last wash; E, eluate; endo, endogenous; epi, episomal.

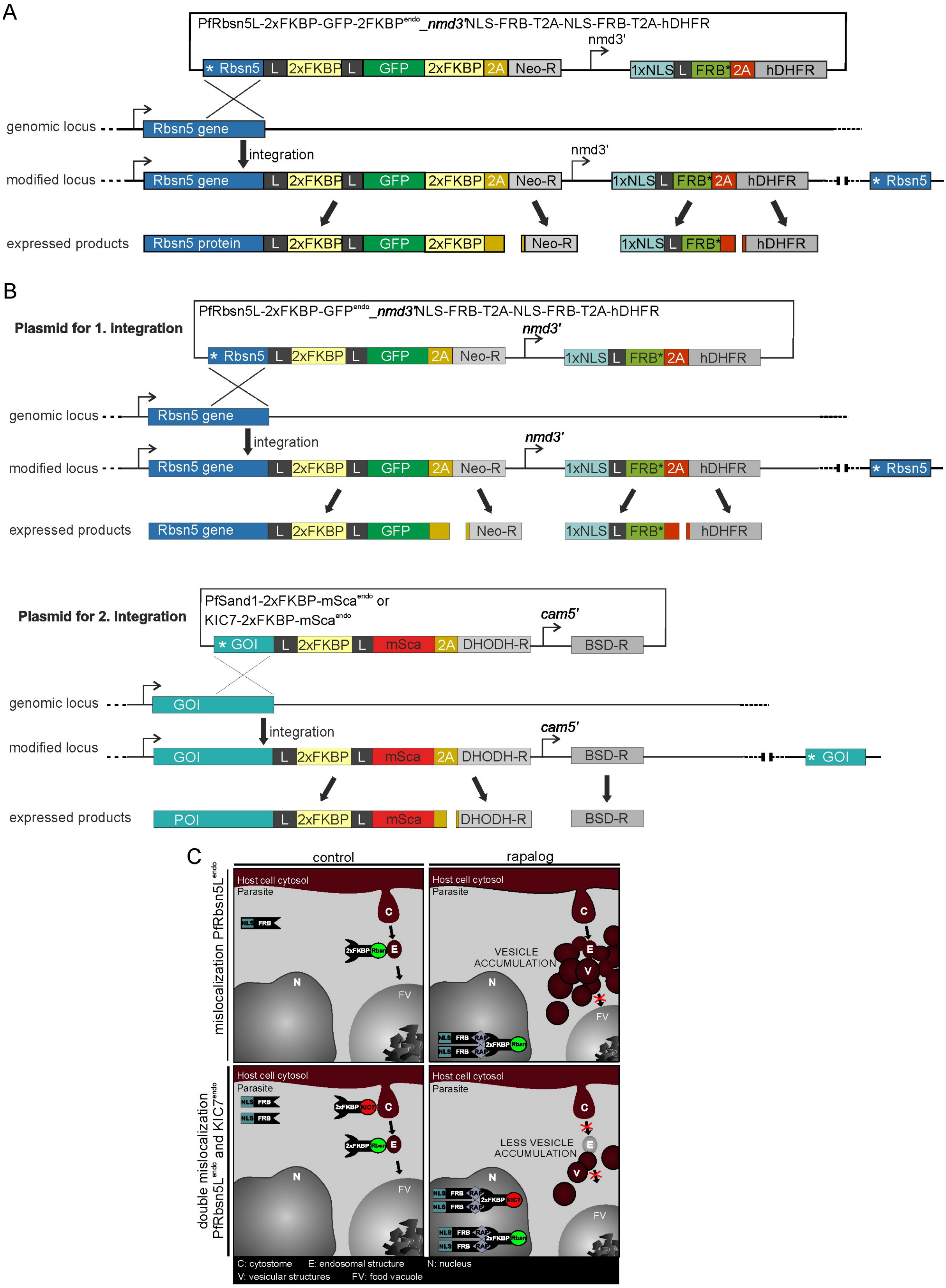


**Figure S9. Modified SLI-sandwich plasmid enabling endogenous tagging and mislocalization of POIs and double conditional inactivation strategy. A,** Schematic of selection linked integration strategy combined with a mislocalization cassette on the same plasmid. **B,** Scheme of a selection linked integration strategy enabling endogenously tagging and simultaneous mislocalization of two POIs in the same parasites. **C,** Schematic illustration of simultaneous double mislocalization of KIC7 (which inhibits endocytosis at the PPM in an early step) and PfRbsn5 (which inhibits transport of HCC-filled vesicles to the food vacuole), showing the expected reduction in vesicle accumulation compared to PfRbsn5 inactivation alone if the two processes are serially linked.

**ADDITIONAL SUPPLEMENTAL FILES**
**Table S1. Primers used for assessing correct genomic integration of plasmids**
**File S1. Plasmids generated for this work.**
