## Supplementary material for "Identification of a Rabenosyn-5 like protein and Rab5b in host cell cytosol uptake reveals conservation of endosomal transport in malaria parasites": Primers to assess plasmid integration into genome

**Table S1**

Oligonucleotides to assess correct integration of plasmids into the genome

**Part A: primer sequences**

| **Primer name** | **Sequence (5'- 3')** |
| --- | --- |
| Rbsn5int_fw | GATTGATAGATCGATATCGATGTGTGG |
| Rbsn5int_rv | GAATAACACATCGACATTTTACATTTTAC |
| pARL55sense | GGAATTGTGAGCGGATAACAATTTCACACAGG |
| GFP85_rv | ACCTTCACCCTCTCCACTGAC |
| Rab5Bint_fw | CGCTAGCTGTTCATATATTATATAAATATAC |
| Rab5Bint_rv | TAGTATATTGATAATATAGAATGACAATG |
| GFP_272_rv | CCTTCGGGCATGGCACTC |
| KIC7int_fw | GATAGAGACAGAGATAAAGATAGAG |
| KIC7int_rv | TACACACATGATATCATACATAAGG |
| Sand1int_fw | TAAGATATTTATTAGATGGAGCAG |
| Sand1int_rv | GATAATATTAATAGTCTATGTAATATC |
| mSca38_rv | CCCTCCATGTGCACCTTAAAACGC |

**Part B: expected length of PCR products**

| **Construct** | **PCR product length** |
| --- | --- |
| - PfRbsn5-2xFKBP-GFP-2xFKBP^endo^  - PfRbsn5-2xFKBP-GFP-2xFKBP^endo^_ndm3'NLS-FRB-T2A-hDHFR  - PfRbsn5-2xFKBP-GFP^endo^ | 5’INT: 1760 bp (Rbsn5int_fw + GFP85_rv);  3’INT: 890 bp (pARL55sense + Rbsn5int_rv)  Ori: 935 bp (Rbsn5int_fw + Rbsn5int_rv) |
| - PfRab5b-2xFKBP-GFP^endo^ | 5’ INT: 1873 bp (Rab5bint_fw + GFP_272_rv)  3’ INT: 896 bp (pARL55sense + Rab5bint_rv)  Ori: 879 bp (Rab5bint_fw + Rab5bint_rv) |
| -KIC7-2xFKBP-mCh^endo^ | 5’INT: 2129 bp (KIC7int_fw + mSca38_rv)  3’INT: 1517 bp (pARL55sense + KIC7int_rv)  Ori: 1479 bp (KIC7int_fw + KIC7int_rv) |
| - Sand1-2xFKBP-mSca^endo^ | 5’INT: 2004 bp (Sand1int_fw + mSca38_rv);  3’INT: 1310 bp (pARL55sense + Sand1int rv)  ori: 1290 bp (Sand1int_fw + Sand1int_rv) |
