## Supplementary material for "Identification of a Rabenosyn-5 like protein and Rab5b in host cell cytosol uptake reveals conservation of endosomal transport in malaria parasites": Sequences of plasmids used

**>PfRbsn5-2xFKBP-GFP-2xFKBP (pSLI)**

XXXX = hDHFR resistance

XXXX = Rbsn5 homology region

XXXX = Linker

XXXX = FKBP

XXXX = GFP

XXXX = T2A skip peptide

XXXX = Neomycin resistance

ggatatggcagcttaatgttcgtttttcttatttatatatttataccaattgattgtatttataactgtaaaaatgtgtatgttgtgtgcatatttttttttgtgcatgcacatgcatgtaaatagctaaaattatgaacattttattttttgttcagaaaaaaaaaactttacacacataaaatggctagtatgaatagccatattttatataaattaaatcctatgaatttatgaccatattaaaaatttagatatttatggaacataatatgtttgaaacaataagacaaaattattattattattattatttttactgttataattatgtgtctccttcaatgattcataaatagttggacttgatttttaaaatgtttataatatgattagcatagttaaataaaaaaagttgaaaaattaaaaaaaaacatataaacacaaatgatggtttttccttcaatttcgatatcaatttatagaaacaaaatatatacttgtataattttatttttttatataaatcattacatatataattatacaatattttttctaagagataattatatattaatatatataaaaaaaggtgttttttttttttttttttatttttatttttattttatggtaatattttattttccttattttataaattatattagtttatatgtgattaattttatatattatcaatttatatatttttaaatgcttacttaattatctttttttttttttttttttttttttcccctctttttatattaatttatttttgaaaaaattgatatatatatatatatataatatatatatatacatgtagtagtattaaacaatgtataatatatataaataatatatttatatatttcatttcaattttaattttttttggttttttttttttttctttttgtcatatttaaaaaaaattatattcatataagttatgcattttttataaacattattcaatatatgtataatataatatatatatatatattaatgtattattccaatgtgcatgataaaagaaaaaaataatatttataaaaaaaaagaaaaataaaacaaaaaaagaaaaaaaaaaaaaaaaaaaaaaaaatacaaaaataaataatataatttataattatatattcttgtcacaataaaaatatatatatatatatatatatttataatatgtatattttaaactagaaaaggaataactaatattttatttattatcattcaagatttatattttataataataaatacctaatagaaatatatcaggatccatgcatggttcgctaaactgcatcgtcgctgtgtcccagaacatgggcatcggcaagaacggggactacccctggccaccgctcaggaacgaatttagatatttccagagaatgaccacaacctcttcagtagaaggtaaacagaatctggtgattatgggtaagaagacctggttctccattcctgagaagaatcgacctttaaagggtagaattaatttagttctcagcagagaactcaaggaacctccacaaggagctcattttctttccagaagtctagatgatgccttaaaacttactgaacaaccagaattagcaaataaagtagacatggtctggatagttggtggcagttctgtttataaggaagccatgaatcacccaggccatcttaaactatttgtgacaaggatcatgcaagactttgaaagtgacacgttttttccagaaattgatttggagaaatataaacttctgccagaatacccaggtgttctctctgatgtccaggaggagaaaggcattaagtacaaatttgaagtatatgagaagaatgattaagcttatttaataatagattaaaaatattataaaaataaaaacataaacacagaaattacaaaaaaaatacatatgaattttttttttgtaatcttccttataaatatagaataatgaatcatataaaacatatcattattcatttatttacatttaaaattattgtttcagtatctttaatttattatgtatatataaaaataacttacaattttattaataaacaatatatgtttattaattcatgttttgtaatttatgggatagcgattttttttactgtctgtatttttcttttttaattatgttttaattgtattttatttttattattgttctttttatagtattattttaaaacaaaatgtattttctaagaacttataataataataaatataaattttaataaaaattatatttatcttttacaatatgaacataaagtacaacattaatatatagcttttaatatttttattcctaatcatgtaaatcttaaatttttctttttaaacatatgttaaatatttatttctcattatatataagaacatatttattaaatctagaattctatagtgagtcgtattacaattcactggccgtcgttttacaacgtcgtgactgggaaaaccctggcgttacccaacttaatcgccttgcagcacatccccctttcgccagctggcgtaatagcgaagaggcccgcaccgatcgcccttcccaacagttgcgcagcctgaatggcgaatggcgcctgatgcggtattttctccttacgcatctgtgcggtatttcacaccgcatatggtgcactctcagtacaatctgctctgatgccgcatagttaagccagccccgacacccgccaacacccgctgacgcgccctgacgggcttgtctgctcccggcatccgcttacagacaagctgtgaccgtctccgggagctgcatgtgtcagaggttttcaccgtcatcaccgaaacgcgcgagacgaaagggcctcgtgatacgcctatttttataggttaatgtcatgataataatggtttcttagacgtcaggtggcacttttcggggaaatgtgcgcggaacccctatttgtttatttttctaaatacattcaaatatgtatccgctcatgagacaataaccctgataaatgcttcaataatattgaaaaaggaagagtatgagtattcaacatttccgtgtcgcccttattcccttttttgcggcattttgccttcctgtttttgctcacccagaaacgctggtgaaagtaaaagatgctgaagatcagttgggtgcacgagtgggttacatcgaactggatctcaacagcggtaagatccttgagagttttcgccccgaagaacgttttccaatgatgagcacttttaaagttctgctatgtggcgcggtattatcccgtattgacgccgggcaagagcaactcggtcgccgcatacactattctcagaatgacttggttgagtactcaccagtcacagaaaagcatcttacggatggcatgacagtaagagaattatgcagtgctgccataaccatgagtgataacactgcggccaacttacttctgacaacgatcggaggaccgaaggagctaaccgcttttttgcacaacatgggggatcatgtaactcgccttgatcgttgggaaccggagctgaatgaagccataccaaacgacgagcgtgacaccacgatgcctgtagcaatgccaacaacgttgcgcaaactattaactggcgaactacttactctagcttcccggcaacaattaatagactggatggaggcggataaagttgcaggaccacttctgcgctcggcccttccggctggctggtttattgctgataaatctggagccggtgagcgtgggtctcgcggtatcattgcagcactggggccagatggtaagccctcccgtatcgtagttatctacacgacggggagtcaggcaactatggatgaacgaaatagacagatcgctgagataggtgcctcactgattaagcattggtaactgtcagaccaagtttactcatatatactttagattgatttaaaacttcatttttaatttaaaaggatctaggtgaagatcctttttgataatctcatgaccaaaatcccttaacgtgagttttcgttccactgagcgtcagaccccgtagaaaagatcaaaggatcttcttgagatcctttttttctgcgcgtaatctgctgcttgcaaacaaaaaaaccaccgctaccagcggtggtttgtttgccggatcaagagctaccaactctttttccgaaggtaactggcttcagcagagcgcagataccaaatactgtccttctagtgtagccgtagttaggccaccacttcaagaactctgtagcaccgcctacatacctcgctctgctaatcctgttaccagtggctgctgccagtggcgataagtcgtgtcttaccgggttggactcaagacgatagttaccggataaggcgcagcggtcgggctgaacggggggttcgtgcacacagcccagcttggagcgaacgacctacaccgaactgagatacctacagcgtgagctatgagaaagcgccacgcttcccgaagggagaaaggcggacaggtatccggtaagcggcagggtcggaacaggagagcgcacgagggagcttccagggggaaacgcctggtatctttatagtcctgtcgggtttcgccacctctgacttgagcgtcgatttttgtgatgctcgtcaggggggcggagcctatcgaaaaacgccagcaacgcggcctttttacggttcctggccttttgctggccttttgctcacatgttctttcctgcgttatcccctgattctgtggataaccgtattaccgcctttgagtgagctgataccgctcgccgcagccgaacgaccgagcgcagcgagtcagtgagcgaggaagcggaagagcgcccaatacgcaaaccgcctctccccgcgcgttggccgattcattaatgcagctggcacgacaggtttcccgactggaaagcgggcagtgagcgcaacgcaattaatgtgagttagctcactcattaggcaccccaggctttacactttatgcttccggctcgtatgttgtgtggaattgtgagcggataacaatttcacacaggaaacagctatgaccatgattacgccaagctatttaggtgacactatagaatactcgcggccgctaaATTGATCCGGTTAAAAGAGATTTGACAAATCTTTTTTTAGAAAAGCGGGAGATGGCAAAAGAAGAGATAGACCCTCGAAAAGATTGGTTAAAGAGAATATTGTTGAATAATAACACGAACGATTTAAGGATGTTTTTAAAGGCTAGCGAATTAAAAACAAAAGAAGGTGATTATTGTATGACATGCAAAAGTAACGTAAAACAGTTATTATATCTTCATACAAAAAAGAATTATTGTCATTTATGTGAAGAAATCTTTTGTGCTTATTGTGTAAAAAGTATTGATTTTATGAAAGATGAAAAAGAGAAATATATAAAAATAAGATTATGTAGAAATTGTTTTATATATATAAATGAACTAAAATATATAATTAATCCAAATTTATCTGTAGATAGAAAAGCTATAGATTTACAAAATTCATTCAATGATATATCAAATTGTTATACAAACTTATGTAGTAACGTACCTCAATTAAATGGATTAGTATTATTATGCGAAAATAATAAAGAATTTTTAGATAGCTTTAAAAGTGAAATTAAACAGTTGGAAGAAAAGATTCAGCATGATGTTGACTTTTTAAATATCATGAAAAAAAAAAATAATTTTTTATCAGACAATACATTAATACTTAATAAAATGGGAAAAAACTTATTATTATATTTGAAAATTATAAGAAATAAAATTATTCCATCATCTATTGAAGTTCTAAATAAAACAAAGGAATTGCTATATAAGAAAcctaggTCAGGATTGAGATCAAGATCTGCTGCTGCTGGTGCTGGTGGTGCTGCTAGAGCTGCTctgcagAGAGGAGTACAAGTTGAAACAATATCACCAGGAGATGGTCGTACATTTCCAAAAAGAGGTCAAACTTGTGTTGTACATTATACTGGAATGCTTGAAGATGGAAAGAAATTTGATTCATCTCGTGATAGAAATAAACCATTTAAATTTATGCTAGGTAAACAAGAAGTAATACGAGGTTGGGAAGAAGGAGTTGCTCAAATGAGTGTAGGTCAAAGAGCAAAACTTACTATATCTCCAGATTATGCTTATGGTGCAACTGGACATCCAGGTATAATTCCACCTCATGCAACTCTTGTATTTGATGTGGAGCTTCTAAAACTAGAAACTAGAGGTGTTCAGGTTGAAACAATTTCACCTGGAGATGGCAGAACCTTTCCTAAAAGAGGACAGACTTGCGTAGTTCATTATACAGGCATGCTAGAGGATGGTAAGAAATTTGATTCTAGTCGAGATAGAAATAAGCCATTCAAGTTTATGCTAGGTAAACAGGAAGTAATAAGAGGTTGGGAAGAGGGTGTAGCACAGATGTCAGTTGGACAAAGAGCAAAGTTAACAATATCACCAGATTATGCATACGGTGCAACAGGCCATCCTGGCATCATCCCTCCACATGCAACTTTAGTATTCGACGTTGAATTGTTAAAGTTAGAGACAacgcgtGCTAGAGGTGCTGCTGCTGGTGCTGGAGGTGCAGGTAGACGTACGATGAGTAAAGGAGAAGAACTTTTCACTGGAGTTGTCCCAATTCTTGTTGAATTAGATGGTGATGTTAATGGGCACAAATTTTCTGTCAGTGGAGAGGGTGAAGGTGATGCAACATACGGAAAACTTACCCTTAAATTTATTTGCACTACTGGAAAACTACCTGTTCCATGGCCAACACTTGTCACTACTTTCGCGTATGGTCTTCAATGCTTTGCGAGATACCCAGATCATATGAAACAGCATGACTTTTTCAAGAGTGCCATGCCCGAAGGTTATGTACAGGAAAGAACTATATTTTTCAAAGATGACGGGAACTACAAGACACGTGCTGAAGTCAAGTTTGAAGGTGATACCCTTGTTAATAGAATCGAGTTAAAAGGTATTGATTTTAAAGAAGATGGAAACATTCTTGGACACAAATTGGAATACAACTATAACTCACACAATGTATACATCATGGCAGACAAACAAAAGAATGGAATCAAAGTTAACTTCAAAATTAGACACAACATTGAAGATGGAAGCGTTCAACTAGCAGACCATTATCAACAAAATACTCCAATTGGCGATGGCCCTGTCCTTTTACCAGACAACCATTACCTGTCCACACAATCTGCCCTTTCGAAAGATCCCAACGAAAAGAGAGACCACATGGTCCTTCTTGAGTTTGTAACAGCTGCTGGGATTACACATGGCATGGATGAGCTCTACAAAGTCGACGCCAGGGGAGCAGCCGCAGGAGCAGGGGGGGCAGGAAGGCGTGGTGTTCAGGTCGAGACTATTAGCCCTGGAGATGGACGCACGTTTCCTAAGCGTGGACAGACATGCGTAGTTCACTACACAGGTATGTTGGAGGACGGTAAAAAGTTCGACAGCTCACGCGACCGCAATAAACCTTTCAAGTTTATGCTTGGCAAGCAGGAGGTTATTCGTGGATGGGAGGAGGGTGTAGCACAGATGTCTGTTGGACAGCGTGCTAAGTTGACAATTTCACCTGACTATGCTTATGGCGCTACGGGCCATCCCGGGATCATTCCGCCACATGCGACTCTGGTATTCGACGTTGAATTATTAAAGTTAGAGACAgctagaggggccgctgcaggtgctggtggagctggaagaCGTGGAGTACAAGTAGAGACTATCTCTCCAGGTGACGGTCGCACTTTCCCAAAGCGTGGCCAAACCTGTGTTGTACATTACACTGGTATGCTGGAGGATGGGAAAAAGTTCGATTCCAGTCGCGACCGTAACAAACCGTTCAAATTCATGTTGGGAAAGCAGGAAGTGATCCGCGGGTGGGAGGAAGGCGTGGCGCAAATGAGCGTCGGTCAGCGGGCTAAATTGACCATTTCCCCTGACTACGCGTATGGGGCTACTGGGCACCCAGGGATTATTCCGCCTCACGCTACACTTGTGTTTGATGTCGAACTTTTGAAACTGGAAACTGTCGACGGAGAAGGAAGAGGAAGTTTATTAACATGTGGAGATGTAGAAGAAAATCCAGGACCAATGATTGAACAAGATGGATTGCACGCAGGTTCTCCGGCCGCTTGGGTGGAGAGGCTATTCGGCTATGACTGGGCACAACAGACAATCGGCTGCTCTGATGCCGCCGTGTTCCGGCTGTCAGCGCAGGGGCGCCCGGTTCTTTTTGTCAAGACCGACCTGTCCGGTGCCCTGAATGAACTGCAGGACGAGGCAGCGCGGCTATCGTGGCTGGCCACGACGGGCGTTCCTTGCGCAGCTGTGCTCGACGTTGTCACTGAAGCGGGAAGGGACTGGCTGCTATTGGGCGAAGTGCCGGGGCAGGATCTCCTGTCATCTCACCTTGCTCCTGCCGAGAAAGTATCCATCATGGCTGATGCAATGCGGCGGCTGCATACGCTTGATCCGGCTACCTGCCCATTCGACCACCAAGCGAAACATCGCATCGAGCGAGCACGTACTCGGATGGAAGCCGGTCTTGTCGATCAGGATGATCTGGACGAAGAGCATCAGGGGCTCGCGCCAGCCGAACTGTTCGCCAGGCTCAAGGCGCGCATGCCCGACGGCGAGGATCTCGTCGTGACCCATGGCGATGCCTGCTTGCCGAATATCATGGTGGAAAATGGCCGCTTTTCTGGATTCATCGACTGTGGCCGGCTGGGTGTGGCGGACCGCTATCAGGACATAGCGTTGGCTACCCGTGATATTGCTGAAGAGCTTGGCGGCGAATGGGCTGACCGCTTCCTCGTGCTTTACGGTATCGCCGCTCCCGATTCGCAGCGCATCGCCTTCTATCGCCTTCTTGACGAGTTCTTCTAACTCGAG

**>PfRab5b-2xFKBP-GFP (pSLI)**

XXXX = hDHFR

XXXX = Rab5b homology region

XXXX = Linker

XXXX = FKBP

XXXX = GFP

XXXX = T2A skip peptide

XXXX = Neomycin resistance

ggatatggcagcttaatgttcgtttttcttatttatatatttataccaattgattgtatttataactgtaaaaatgtgtatgttgtgtgcatatttttttttgtgcatgcacatgcatgtaaatagctaaaattatgaacattttattttttgttcagaaaaaaaaaactttacacacataaaatggctagtatgaatagccatattttatataaattaaatcctatgaatttatgaccatattaaaaatttagatatttatggaacataatatgtttgaaacaataagacaaaattattattattattattatttttactgttataattatgtgtctccttcaatgattcataaatagttggacttgatttttaaaatgtttataatatgattagcatagttaaataaaaaaagttgaaaaattaaaaaaaaacatataaacacaaatgatggtttttccttcaatttcgatatcaatttatagaaacaaaatatatacttgtataattttatttttttatataaatcattacatatataattatacaatattttttctaagagataattatatattaatatatataaaaaaaggtgttttttttttttttttttatttttatttttattttatggtaatattttattttccttattttataaattatattagtttatatgtgattaattttatatattatcaatttatatatttttaaatgcttacttaattatctttttttttttttttttttttttttcccctctttttatattaatttatttttgaaaaaattgatatatatatatatatataatatatatatatacatgtagtagtattaaacaatgtataatatatataaataatatatttatatatttcatttcaattttaattttttttggttttttttttttttctttttgtcatatttaaaaaaaattatattcatataagttatgcattttttataaacattattcaatatatgtataatataatatatatatatatattaatgtattattccaatgtgcatgataaaagaaaaaaataatatttataaaaaaaaagaaaaataaaacaaaaaaagaaaaaaaaaaaaaaaaaaaaaaaaatacaaaaataaataatataatttataattatatattcttgtcacaataaaaatatatatatatatatatatatttataatatgtatattttaaactagaaaaggaataactaatattttatttattatcattcaagatttatattttataataataaatacctaatagaaatatatcaggatccatgcatggttcgctaaactgcatcgtcgctgtgtcccagaacatgggcatcggcaagaacggggactacccctggccaccgctcaggaacgaatttagatatttccagagaatgaccacaacctcttcagtagaaggtaaacagaatctggtgattatgggtaagaagacctggttctccattcctgagaagaatcgacctttaaagggtagaattaatttagttctcagcagagaactcaaggaacctccacaaggagctcattttctttccagaagtctagatgatgccttaaaacttactgaacaaccagaattagcaaataaagtagacatggtctggatagttggtggcagttctgtttataaggaagccatgaatcacccaggccatcttaaactatttgtgacaaggatcatgcaagactttgaaagtgacacgttttttccagaaattgatttggagaaatataaacttctgccagaatacccaggtgttctctctgatgtccaggaggagaaaggcattaagtacaaatttgaagtatatgagaagaatgattaagcttatttaataatagattaaaaatattataaaaataaaaacataaacacagaaattacaaaaaaaatacatatgaattttttttttgtaatcttccttataaatatagaataatgaatcatataaaacatatcattattcatttatttacatttaaaattattgtttcagtatctttaatttattatgtatatataaaaataacttacaattttattaataaacaatatatgtttattaattcatgttttgtaatttatgggatagcgattttttttactgtctgtatttttcttttttaattatgttttaattgtattttatttttattattgttctttttatagtattattttaaaacaaaatgtattttctaagaacttataataataataaatataaattttaataaaaattatatttatcttttacaatatgaacataaagtacaacattaatatatagcttttaatatttttattcctaatcatgtaaatcttaaatttttctttttaaacatatgttaaatatttatttctcattatatataagaacatatttattaaatctagaattctatagtgagtcgtattacaattcactggccgtcgttttacaacgtcgtgactgggaaaaccctggcgttacccaacttaatcgccttgcagcacatccccctttcgccagctggcgtaatagcgaagaggcccgcaccgatcgcccttcccaacagttgcgcagcctgaatggcgaatggcgcctgatgcggtattttctccttacgcatctgtgcggtatttcacaccgcatatggtgcactctcagtacaatctgctctgatgccgcatagttaagccagccccgacacccgccaacacccgctgacgcgccctgacgggcttgtctgctcccggcatccgcttacagacaagctgtgaccgtctccgggagctgcatgtgtcagaggttttcaccgtcatcaccgaaacgcgcgagacgaaagggcctcgtgatacgcctatttttataggttaatgtcatgataataatggtttcttagacgtcaggtggcacttttcggggaaatgtgcgcggaacccctatttgtttatttttctaaatacattcaaatatgtatccgctcatgagacaataaccctgataaatgcttcaataatattgaaaaaggaagagtatgagtattcaacatttccgtgtcgcccttattcccttttttgcggcattttgccttcctgtttttgctcacccagaaacgctggtgaaagtaaaagatgctgaagatcagttgggtgcacgagtgggttacatcgaactggatctcaacagcggtaagatccttgagagttttcgccccgaagaacgttttccaatgatgagcacttttaaagttctgctatgtggcgcggtattatcccgtattgacgccgggcaagagcaactcggtcgccgcatacactattctcagaatgacttggttgagtactcaccagtcacagaaaagcatcttacggatggcatgacagtaagagaattatgcagtgctgccataaccatgagtgataacactgcggccaacttacttctgacaacgatcggaggaccgaaggagctaaccgcttttttgcacaacatgggggatcatgtaactcgccttgatcgttgggaaccggagctgaatgaagccataccaaacgacgagcgtgacaccacgatgcctgtagcaatgccaacaacgttgcgcaaactattaactggcgaactacttactctagcttcccggcaacaattaatagactggatggaggcggataaagttgcaggaccacttctgcgctcggcccttccggctggctggtttattgctgataaatctggagccggtgagcgtgggtctcgcggtatcattgcagcactggggccagatggtaagccctcccgtatcgtagttatctacacgacggggagtcaggcaactatggatgaacgaaatagacagatcgctgagataggtgcctcactgattaagcattggtaactgtcagaccaagtttactcatatatactttagattgatttaaaacttcatttttaatttaaaaggatctaggtgaagatcctttttgataatctcatgaccaaaatcccttaacgtgagttttcgttccactgagcgtcagaccccgtagaaaagatcaaaggatcttcttgagatcctttttttctgcgcgtaatctgctgcttgcaaacaaaaaaaccaccgctaccagcggtggtttgtttgccggatcaagagctaccaactctttttccgaaggtaactggcttcagcagagcgcagataccaaatactgtccttctagtgtagccgtagttaggccaccacttcaagaactctgtagcaccgcctacatacctcgctctgctaatcctgttaccagtggctgctgccagtggcgataagtcgtgtcttaccgggttggactcaagacgatagttaccggataaggcgcagcggtcgggctgaacggggggttcgtgcacacagcccagcttggagcgaacgacctacaccgaactgagatacctacagcgtgagctatgagaaagcgccacgcttcccgaagggagaaaggcggacaggtatccggtaagcggcagggtcggaacaggagagcgcacgagggagcttccagggggaaacgcctggtatctttatagtcctgtcgggtttcgccacctctgacttgagcgtcgatttttgtgatgctcgtcaggggggcggagcctatcgaaaaacgccagcaacgcggcctttttacggttcctggccttttgctggccttttgctcacatgttctttcctgcgttatcccctgattctgtggataaccgtattaccgcctttgagtgagctgataccgctcgccgcagccgaacgaccgagcgcagcgagtcagtgagcgaggaagcggaagagcgcccaatacgcaaaccgcctctccccgcgcgttggccgattcattaatgcagctggcacgacaggtttcccgactggaaagcgggcagtgagcgcaacgcaattaatgtgagttagctcactcattaggcaccccaggctttacactttatgcttccggctcgtatgttgtgtggaattgtgagcggataacaatttcacacaggaaacagctatgaccatgattacgccaagctatttaggtgacactatagaatactcgcggccgctaaTAATGTCGAATCCTTTGATTCTTTAAAATACTGGATAAACGAAATAAAATCAAATGGTCCAAGAAATTGTTGTATCATGGTAGTTGCCAATAAAAAGGACTTACCCCAAAAACTGAATTCAGAGgtaatcatgaggacacattaaaaagtttatatattcatatagataagaaaaatgatatatatatatatatatatatatatatatatatatatttatatttatatatttatatatgtatgtcaatttatttaactttttattttattagATGGTCATGAAGTTTTGTGAACAAGAAAATGTATCATTTATTGAATGCTCAGCTAAGgtataaatattcaaaaaaaatatattaataaatttataacatattataagaatagcaacatataagagttttacaattatttgattccctctctataaatatatatatatatatatatatatatatttttttagACAGGAGAGAATATTACTACTTTATTTGAAAAATTAGgttagattatttataatttttacatatttccaatgaaaataattgaataattaaataaaaaaataaaatctataatactacaataatatcttatttatattttatattatattatattatattatattatattatattatattatattatattatattatattatattatattattttttattttttgttttacagCTAGTCGCATTTATTCCCGATTTAAAGAAGTTTTATATTATAACAATCCTcctaggTCAGGATTGAGATCAAGATCTGCTGCTGCTGGTGCTGGTGGTGCTGCTAGAGCTGCTctgcagAGAGGAGTACAAGTTGAAACAATATCACCAGGAGATGGTCGTACATTTCCAAAAAGAGGTCAAACTTGTGTTGTACATTATACTGGAATGCTTGAAGATGGAAAGAAATTTGATTCATCTCGTGATAGAAATAAACCATTTAAATTTATGCTAGGTAAACAAGAAGTAATACGAGGTTGGGAAGAAGGAGTTGCTCAAATGAGTGTAGGTCAAAGAGCAAAACTTACTATATCTCCAGATTATGCTTATGGTGCAACTGGACATCCAGGTATAATTCCACCTCATGCAACTCTTGTATTTGATGTGGAGCTTCTAAAACTAGAAACTAGAGGTGTTCAGGTTGAAACAATTTCACCTGGAGATGGCAGAACCTTTCCTAAAAGAGGACAGACTTGCGTAGTTCATTATACAGGCATGCTAGAGGATGGTAAGAAATTTGATTCTAGTCGAGATAGAAATAAGCCATTCAAGTTTATGCTAGGTAAACAGGAAGTAATAAGAGGTTGGGAAGAGGGTGTAGCACAGATGTCAGTTGGACAAAGAGCAAAGTTAACAATATCACCAGATTATGCATACGGTGCAACAGGCCATCCTGGCATCATCCCTCCACATGCAACTTTAGTATTCGACGTTGAATTGTTAAAGTTAGAGACAacgcgtGCTAGAGGTGCTGCTGCTGGTGCTGGAGGTGCAGGTAGACGTACGATGAGTAAAGGAGAAGAACTTTTCACTGGAGTTGTCCCAATTCTTGTTGAATTAGATGGTGATGTTAATGGGCACAAATTTTCTGTCAGTGGAGAGGGTGAAGGTGATGCAACATACGGAAAACTTACCCTTAAATTTATTTGCACTACTGGAAAACTACCTGTTCCATGGCCAACACTTGTCACTACTTTCGCGTATGGTCTTCAATGCTTTGCGAGATACCCAGATCATATGAAACAGCATGACTTTTTCAAGAGTGCCATGCCCGAAGGTTATGTACAGGAAAGAACTATATTTTTCAAAGATGACGGGAACTACAAGACACGTGCTGAAGTCAAGTTTGAAGGTGATACCCTTGTTAATAGAATCGAGTTAAAAGGTATTGATTTTAAAGAAGATGGAAACATTCTTGGACACAAATTGGAATACAACTATAACTCACACAATGTATACATCATGGCAGACAAACAAAAGAATGGAATCAAAGTTAACTTCAAAATTAGACACAACATTGAAGATGGAAGCGTTCAACTAGCAGACCATTATCAACAAAATACTCCAATTGGCGATGGCCCTGTCCTTTTACCAGACAACCATTACCTGTCCACACAATCTGCCCTTTCGAAAGATCCCAACGAAAAGAGAGACCACATGGTCCTTCTTGAGTTTGTAACAGCTGCTGGGATTACACATGGCATGGATGAGCTCTACAAAGTCGACGGAGAAGGAAGAGGAAGTTTATTAACATGTGGAGATGTAGAAGAAAATCCAGGACCAATGATTGAACAAGATGGATTGCACGCAGGTTCTCCGGCCGCTTGGGTGGAGAGGCTATTCGGCTATGACTGGGCACAACAGACAATCGGCTGCTCTGATGCCGCCGTGTTCCGGCTGTCAGCGCAGGGGCGCCCGGTTCTTTTTGTCAAGACCGACCTGTCCGGTGCCCTGAATGAACTGCAGGACGAGGCAGCGCGGCTATCGTGGCTGGCCACGACGGGCGTTCCTTGCGCAGCTGTGCTCGACGTTGTCACTGAAGCGGGAAGGGACTGGCTGCTATTGGGCGAAGTGCCGGGGCAGGATCTCCTGTCATCTCACCTTGCTCCTGCCGAGAAAGTATCCATCATGGCTGATGCAATGCGGCGGCTGCATACGCTTGATCCGGCTACCTGCCCATTCGACCACCAAGCGAAACATCGCATCGAGCGAGCACGTACTCGGATGGAAGCCGGTCTTGTCGATCAGGATGATCTGGACGAAGAGCATCAGGGGCTCGCGCCAGCCGAACTGTTCGCCAGGCTCAAGGCGCGCATGCCCGACGGCGAGGATCTCGTCGTGACCCATGGCGATGCCTGCTTGCCGAATATCATGGTGGAAAATGGCCGCTTTTCTGGATTCATCGACTGTGGCCGGCTGGGTGTGGCGGACCGCTATCAGGACATAGCGTTGGCTACCCGTGATATTGCTGAAGAGCTTGGCGGCGAATGGGCTGACCGCTTCCTCGTGCTTTACGGTATCGCCGCTCCCGATTCGCAGCGCATCGCCTTCTATCGCCTTCTTGACGAGTTCTTCTAACTCGAG

**>PfRbsn5-2x2/NLS (pSLI)**

XXXX = hDHFR resistance

XXXX = Rbsn5 homology region

XXXX = Linker

XXXX = FKBP

XXXX = GFP

XXXX = T2A skip peptide

XXXX = Neomycin resistance

XXXX = FRB

XXXX = NLS

XXXX = *nmd3’* promotor

ggatatggcagcttaatgttcgtttttcttatttatatatttataccaattgattgtatttataactgtaaaaatgtgtatgttgtgtgcatatttttttttgtgcatgcacatgcatgtaaatagctaaaattatgaacattttattttttgttcagaaaaaaaaaactttacacacataaaatggctagtatgaatagccatattttatataaattaaatcctatgaatttatgaccatattaaaaatttagatatttatggaacataatatgtttgaaacaataagacaaaattattattattattattatttttactgttataattatgtgtctccttcaatgattcataaatagttggacttgatttttaaaatgtttataatatgattagcatagttaaataaaaaaagttgaaaaattaaaaaaaaacatataaacacaaatgatggtttttccttcaatttcgatatcaatttatagaaacaaaatatatacttgtataattttatttttttatataaatcattacatatataattatacaatattttttctaagagataattatatattaatatatataaaaaaaggtgttttttttttttttttttatttttatttttattttatggtaatattttattttccttattttataaattatattagtttatatgtgattaattttatatattatcaatttatatatttttaaatgcttacttaattatctttttttttttttttttttttttttcccctctttttatattaatttatttttgaaaaaattgatatatatataGTTGAAATATAAATTTCAAAAAAAATGATCACAAAATATACACTTAAATATAGGTACAATAAAAAAAAAAAATAAAAATATAATTACAAGATAATATTTTTTCCTGCTATCAAATTTTTATATATTCTCCTCAAAGAAAAAATATAAATAAATGAAGTAAATTAAAAAAAAAATTTTCTTTTTCTTCTTCTTTTGTAATTCCTTATTTATACATATTTTACTATATTTTCATAAAAATAAATTGTCATATTATATAAATATATATACCAAACCATAATTATATAGCCCTCAACATATTTTTATGATGTTTTTTCTTTTTAAATGTGATACGTAATTAAATATAATAATATATATTAAATATTATATTTTGTAATATTTACTTTCATGAGGTTTAATAAATTATAAAGAGAGAATAAAAAAAAAAAAAAAAAAAATTTATCCATATACAAATTATTAATTTATTTTTATTTTTTATTTCCCTTTGTATATATTATAAAAAAATAATCTACATAATTTTATATGATGATAATTACAATATAATATTTATAATATATATTATTTGTTAAGAGAAAAAAAAAATAAATATATACCTTCTTTTTGAGAATTGAATAAATTGTTTAATATATATATATATATATATAAATATATTATACATTTATGGTGAAAAAAAATGTTGTATTTAATTAGATTTAATATATATATATAAAAAATAGCGTATTTAAAATAATATATATATATATATATTATTATTATTACAAAATGACGGATATTATAAAAGTATATATCTATATATATGTATATATATATAATATTATTTTACTATATATATATAATATATAAATGTAAATGCATATAGTATCTATGTATATTATATATATATATAATATTAATTACATATCTAAGTTTTCTTTTCTTTTCTTTTTTTTTTTTTTTTTTTTTATTTTTTATAGAGAGCCCGTTATATATTATATATTAAAATTTTTTATAATAGCATTTATATACAATATTTTATTAACTAAAAGAAAAAAAAAAAAAAAAAAAAAAAAAAAACGAGGAAATTTATATTTCTTTAACAACATTTTAATTAAAATCATGATAATACAATTTCAAATCATTTTGTAATTATATAAATAAATATATATATATATATATATATATATATGTACTTTTAAATTAGGAATATTCTCATTTATAAATATATCTTATTTTTTAAATTGGTATAAAAAAAAAAAAAAAAATAAGAAACCGTTGATTAAATAATACATATATAATATAAATATATTTTATAAATATATATTTATATATATATATATATATATTTATAACGTATATCATTTTAAAGATAAACTAGTatggcaccaaaaaaaaaaagaaaagttacgcgtgatccaacaagaagtgcaaatagtggagcaggagcaggagcaggagcaatattaagtagagctagcatggcttctagaatcctctggcatgagatGtggcatgaaggcctggaagaggcatctcgtttgtactttggggaaaggaacgtgaaaggCatgtttgaggtgctggagcccttgcatgctatgatggaacggggcccccagactctgaaGgaaacatcctttaatcaggcctatggtcgagatttaatggaggcccaagagtggtgcagGaagtacatgaaatcagggaatgtcaaggacctcctccaagcctgggacctctattatcaTgtgttccgacgaatctcaaagGGTGAGGGTCGTGGTTCACTTCTTACTTGCGGTGACGTTGAGGAGAACCCTGGTCCTGGTACCggatcccatgcatggttcgctaaactgcatcgtcgctgtgtcccagaacatgggcatcggcaagaacggggactacccctggccaccgctcaggaacgaatttagatatttccagagaatgaccacaacctcttcagtagaaggtaaacagaatctggtgattatgggtaagaagacctggttctccattcctgagaagaatcgacctttaaagggtagaattaatttagttctcagcagagaactcaaggaacctccacaaggagctcattttctttccagaagtctagatgatgccttaaaacttactgaacaaccagaattagcaaataaagtagacatggtctggatagttggtggcagttctgtttataaggaagccatgaatcacccaggccatcttaaactatttgtgacaaggatcatgcaagactttgaaagtgacacgttttttccagaaattgatttggagaaatataaacttctgccagaatacccaggtgttctctctgatgtccaggaggagaaaggcattaagtacaaatttgaagtatatgagaagaatgattaagcttatttaataatagattaaaaatattataaaaataaaaacataaacacagaaattacaaaaaaaatacatatgaattttttttttgtaatcttccttataaatatagaataatgaatcatataaaacatatcattattcatttatttacatttaaaattattgtttcagtatctttaatttattatgtatatataaaaataacttacaattttattaataaacaatatatgtttattaattcatgttttgtaatttatgggatagcgattttttttactgtctgtatttttcttttttaattatgttttaattgtattttatttttattattgttctttttatagtattattttaaaacaaaatgtattttctaagaacttataataataataaatataaattttaataaaaattatatttatcttttacaatatgaacataaagtacaacattaatatatagcttttaatatttttattcctaatcatgtaaatcttaaatttttctttttaaacatatgttaaatatttatttctcattatatataagaacatatttattaaatctagaattctatagtgagtcgtattacaattcactggccgtcgttttacaacgtcgtgactgggaaaaccctggcgttacccaacttaatcgccttgcagcacatccccctttcgccagctggcgtaatagcgaagaggcccgcaccgatcgcccttcccaacagttgcgcagcctgaatggcgaatggcgcctgatgcggtattttctccttacgcatctgtgcggtatttcacaccgcatatggtgcactctcagtacaatctgctctgatgccgcatagttaagccagccccgacacccgccaacacccgctgacgcgccctgacgggcttgtctgctcccggcatccgcttacagacaagctgtgaccgtctccgggagctgcatgtgtcagaggttttcaccgtcatcaccgaaacgcgcgagacgaaagggcctcgtgatacgcctatttttataggttaatgtcatgataataatggtttcttagacgtcaggtggcacttttcggggaaatgtgcgcggaacccctatttgtttatttttctaaatacattcaaatatgtatccgctcatgagacaataaccctgataaatgcttcaataatattgaaaaaggaagagtatgagtattcaacatttccgtgtcgcccttattcccttttttgcggcattttgccttcctgtttttgctcacccagaaacgctggtgaaagtaaaagatgctgaagatcagttgggtgcacgagtgggttacatcgaactggatctcaacagcggtaagatccttgagagttttcgccccgaagaacgttttccaatgatgagcacttttaaagttctgctatgtggcgcggtattatcccgtattgacgccgggcaagagcaactcggtcgccgcatacactattctcagaatgacttggttgagtactcaccagtcacagaaaagcatcttacggatggcatgacagtaagagaattatgcagtgctgccataaccatgagtgataacactgcggccaacttacttctgacaacgatcggaggaccgaaggagctaaccgcttttttgcacaacatgggggatcatgtaactcgccttgatcgttgggaaccggagctgaatgaagccataccaaacgacgagcgtgacaccacgatgcctgtagcaatgccaacaacgttgcgcaaactattaactggcgaactacttactctagcttcccggcaacaattaatagactggatggaggcggataaagttgcaggaccacttctgcgctcggcccttccggctggctggtttattgctgataaatctggagccggtgagcgtgggtctcgcggtatcattgcagcactggggccagatggtaagccctcccgtatcgtagttatctacacgacggggagtcaggcaactatggatgaacgaaatagacagatcgctgagataggtgcctcactgattaagcattggtaactgtcagaccaagtttactcatatatactttagattgatttaaaacttcatttttaatttaaaaggatctaggtgaagatcctttttgataatctcatgaccaaaatcccttaacgtgagttttcgttccactgagcgtcagaccccgtagaaaagatcaaaggatcttcttgagatcctttttttctgcgcgtaatctgctgcttgcaaacaaaaaaaccaccgctaccagcggtggtttgtttgccggatcaagagctaccaactctttttccgaaggtaactggcttcagcagagcgcagataccaaatactgtccttctagtgtagccgtagttaggccaccacttcaagaactctgtagcaccgcctacatacctcgctctgctaatcctgttaccagtggctgctgccagtggcgataagtcgtgtcttaccgggttggactcaagacgatagttaccggataaggcgcagcggtcgggctgaacggggggttcgtgcacacagcccagcttggagcgaacgacctacaccgaactgagatacctacagcgtgagctatgagaaagcgccacgcttcccgaagggagaaaggcggacaggtatccggtaagcggcagggtcggaacaggagagcgcacgagggagcttccagggggaaacgcctggtatctttatagtcctgtcgggtttcgccacctctgacttgagcgtcgatttttgtgatgctcgtcaggggggcggagcctatcgaaaaacgccagcaacgcggcctttttacggttcctggccttttgctggccttttgctcacatgttctttcctgcgttatcccctgattctgtggataaccgtattaccgcctttgagtgagctgataccgctcgccgcagccgaacgaccgagcgcagcgagtcagtgagcgaggaagcggaagagcgcccaatacgcaaaccgcctctccccgcgcgttggccgattcattaatgcagctggcacgacaggtttcccgactggaaagcgggcagtgagcgcaacgcaattaatgtgagttagctcactcattaggcaccccaggctttacactttatgcttccggctcgtatgttgtgtggaattgtgagcggataacaatttcacacaggaaacagctatgaccatgattacgccaagctatttaggtgacactatagaatactcgcggccgctaaATTGATCCGGTTAAAAGAGATTTGACAAATCTTTTTTTAGAAAAGCGGGAGATGGCAAAAGAAGAGATAGACCCTCGAAAAGATTGGTTAAAGAGAATATTGTTGAATAATAACACGAACGATTTAAGGATGTTTTTAAAGGCTAGCGAATTAAAAACAAAAGAAGGTGATTATTGTATGACATGCAAAAGTAACGTAAAACAGTTATTATATCTTCATACAAAAAAGAATTATTGTCATTTATGTGAAGAAATCTTTTGTGCTTATTGTGTAAAAAGTATTGATTTTATGAAAGATGAAAAAGAGAAATATATAAAAATAAGATTATGTAGAAATTGTTTTATATATATAAATGAACTAAAATATATAATTAATCCAAATTTATCTGTAGATAGAAAAGCTATAGATTTACAAAATTCATTCAATGATATATCAAATTGTTATACAAACTTATGTAGTAACGTACCTCAATTAAATGGATTAGTATTATTATGCGAAAATAATAAAGAATTTTTAGATAGCTTTAAAAGTGAAATTAAACAGTTGGAAGAAAAGATTCAGCATGATGTTGACTTTTTAAATATCATGAAAAAAAAAAATAATTTTTTATCAGACAATACATTAATACTTAATAAAATGGGAAAAAACTTATTATTATATTTGAAAATTATAAGAAATAAAATTATTCCATCATCTATTGAAGTTCTAAATAAAACAAAGGAATTGCTATATAAGAAAcctaggTCAGGATTGAGATCAAGATCTGCTGCTGCTGGTGCTGGTGGTGCTGCTAGAGCTGCTctgcagTAGTATTCGACGTTGAATTGTTAAAGTTAGAGACATGGAAAGAAATTTGATTCATCTCGTGATAGAAATAAACCATTTAAATTTATGCTAGGTAAACAAGAAGTAATACGAGGTTGGGAAGAAGGAGTTGCTCAAATGAGTGTAGGTCAAAGAGCAAAACTTACTATATCTCCAGATTATGCTTATGGTGCAACTGGACATCCAGGTATAATTCCACCTCATGCAACTCTTGTATTTGATGTGGAGCTTCTAAAACTAGAAACTAGAGGTGTTCAGGTTGAAACAATTTCACCTGGAGATGGCAGAACCTTTCCTAAAAGAGGACAGACTTGCGTAGTTCATTATACAGGCATGCTAGAGGATGGTAAGAAATTTGATTCTAGTCGAGATAGAAATAAGCCATTCAAGTTTATGCTAGGTAAACAGGAAGTAATAAGAGGTTGGGAAGAGGGTGTAGCACAGATGTCAGTTGGACAAAGAGCAAAGTTAACAATATCACCAGATTATGCATACGGTGCAACAGGCCATCCTGGCATCATCCCTCCACATGCAACTTTAGTATTCGACGTTGAATTGTTAAAGTTAGAGACAacgcgtGCTAGAGGTGCTGCTGCTGGTGCTGGAGGTGCAGGTAGACGTACGATGAGTAAAGGAGAAGAACTTTTCACTGGAGTTGTCCCAATTCTTGTTGAATTAGATGGTGATGTTAATGGGCACAAATTTTCTGTCAGTGGAGAGGGTGAAGGTGATGCAACATACGGAAAACTTACCCTTAAATTTATTTGCACTACTGGAAAACTACCTGTTCCATGGCCAACACTTGTCACTACTTTCGCGTATGGTCTTCAATGCTTTGCGAGATACCCAGATCATATGAAACAGCATGACTTTTTCAAGAGTGCCATGCCCGAAGGTTATGTACAGGAAAGAACTATATTTTTCAAAGATGACGGGAACTACAAGACACGTGCTGAAGTCAAGTTTGAAGGTGATACCCTTGTTAATAGAATCGAGTTAAAAGGTATTGATTTTAAAGAAGATGGAAACATTCTTGGACACAAATTGGAATACAACTATAACTCACACAATGTATACATCATGGCAGACAAACAAAAGAATGGAATCAAAGTTAACTTCAAAATTAGACACAACATTGAAGATGGAAGCGTTCAACTAGCAGACCATTATCAACAAAATACTCCAATTGGCGATGGCCCTGTCCTTTTACCAGACAACCATTACCTGTCCACACAATCTGCCCTTTCGAAAGATCCCAACGAAAAGAGAGACCACATGGTCCTTCTTGAGTTTGTAACAGCTGCTGGGATTACACATGGCATGGATGAGCTCTACAAAGTCGACGCCAGGGGAGCAGCCGCAGGAGCAGGGGGGGCAGGAAGGCGTGGTGTTCAGGTCGAGACTATTAGCCCTGGAGATGGACGCACGTTTCCTAAGCGTGGACAGACATGCGTAGTTCACTACACAGGTATGTTGGAGGACGGTAAAAAGTTCGACAGCTCACGCGACCGCAATAAACCTTTCAAGTTTATGCTTGGCAAGCAGGAGGTTATTCGTGGATGGGAGGAGGGTGTAGCACAGATGTCTGTTGGACAGCGTGCTAAGTTGACAATTTCACCTGACTATGCTTATGGCGCTACGGGCCATCCCGGGATCATTCCGCCACATGCGACTCTGGTATTCGACGTTGAATTATTAAAGTTAGAGACAgctagaggggccgctgcaggtgctggtggagctggaagaCGTGGAGTACAAGTAGAGACTATCTCTCCAGGTGACGGTCGCACTTTCCCAAAGCGTGGCCAAACCTGTGTTGTACATTACACTGGTATGCTGGAGGATGGGAAAAAGTTCGATTCCAGTCGCGACCGTAACAAACCGTTCAAATTCATGTTGGGAAAGCAGGAAGTGATCCGCGGGTGGGAGGAAGGCGTGGCGCAAATGAGCGTCGGTCAGCGGGCTAAATTGACCATTTCCCCTGACTACGCGTATGGGGCTACTGGGCACCCAGGGATTATTCCGCCTCACGCTACACTTGTGTTTGATGTCGAACTTTTGAAACTGGAAACTGTCGACGGAGAAGGAAGAGGAAGTTTATTAACATGTGGAGATGTAGAAGAAAATCCAGGACCAATGATTGAACAAGATGGATTGCACGCAGGTTCTCCGGCCGCTTGGGTGGAGAGGCTATTCGGCTATGACTGGGCACAACAGACAATCGGCTGCTCTGATGCCGCCGTGTTCCGGCTGTCAGCGCAGGGGCGCCCGGTTCTTTTTGTCAAGACCGACCTGTCCGGTGCCCTGAATGAACTGCAGGACGAGGCAGCGCGGCTATCGTGGCTGGCCACGACGGGCGTTCCTTGCGCAGCTGTGCTCGACGTTGTCACTGAAGCGGGAAGGGACTGGCTGCTATTGGGCGAAGTGCCGGGGCAGGATCTCCTGTCATCTCACCTTGCTCCTGCCGAGAAAGTATCCATCATGGCTGATGCAATGCGGCGGCTGCATACGCTTGATCCGGCTACCTGCCCATTCGACCACCAAGCGAAACATCGCATCGAGCGAGCACGTACTCGGATGGAAGCCGGTCTTGTCGATCAGGATGATCTGGACGAAGAGCATCAGGGGCTCGCGCCAGCCGAACTGTTCGCCAGGCTCAAGGCGCGCATGCCCGACGGCGAGGATCTCGTCGTGACCCATGGCGATGCCTGCTTGCCGAATATCATGGTGGAAAATGGCCGCTTTTCTGGATTCATCGACTGTGGCCGGCTGGGTGTGGCGGACCGCTATCAGGACATAGCGTTGGCTACCCGTGATATTGCTGAAGAGCTTGGCGGCGAATGGGCTGACCGCTTCCTCGTGCTTTACGGTATCGCCGCTCCCGATTCGCAGCGCATCGCCTTCTATCGCCTTCTTGACGAGTTCTTCTAACTCGAG

**>PfRbsn5-2x/NLS (pSLI)**

XXXX = hDHFR resistance

XXXX = Rbsn5 homology region

XXXX = Linker

XXXX = FKBP

XXXX = GFP

XXXX = T2A skip peptide

XXXX = Neomycin resistance

XXXX = FRB

XXXX = NLS

XXXX = *nmd3’* promotor

ggatatggcagcttaatgttcgtttttcttatttatatatttataccaattgattgtatttataactgtaaaaatgtgtatgttgtgtgcatatttttttttgtgcatgcacatgcatgtaaatagctaaaattatgaacattttattttttgttcagaaaaaaaaaactttacacacataaaatggctagtatgaatagccatattttatataaattaaatcctatgaatttatgaccatattaaaaatttagatatttatggaacataatatgtttgaaacaataagacaaaattattattattattattatttttactgttataattatgtgtctccttcaatgattcataaatagttggacttgatttttaaaatgtttataatatgattagcatagttaaataaaaaaagttgaaaaattaaaaaaaaacatataaacacaaatgatggtttttccttcaatttcgatatcaatttatagaaacaaaatatatacttgtataattttatttttttatataaatcattacatatataattatacaatattttttctaagagataattatatattaatatatataaaaaaaggtgttttttttttttttttttatttttatttttattttatggtaatattttattttccttattttataaattatattagtttatatgtgattaattttatatattatcaatttatatatttttaaatgcttacttaattatctttttttttttttttttttttttttcccctctttttatattaatttatttttgaaaaaattgatatatatataGTTGAAATATAAATTTCAAAAAAAATGATCACAAAATATACACTTAAATATAGGTACAATAAAAAAAAAAAATAAAAATATAATTACAAGATAATATTTTTTCCTGCTATCAAATTTTTATATATTCTCCTCAAAGAAAAAATATAAATAAATGAAGTAAATTAAAAAAAAAATTTTCTTTTTCTTCTTCTTTTGTAATTCCTTATTTATACATATTTTACTATATTTTCATAAAAATAAATTGTCATATTATATAAATATATATACCAAACCATAATTATATAGCCCTCAACATATTTTTATGATGTTTTTTCTTTTTAAATGTGATACGTAATTAAATATAATAATATATATTAAATATTATATTTTGTAATATTTACTTTCATGAGGTTTAATAAATTATAAAGAGAGAATAAAAAAAAAAAAAAAAAAAATTTATCCATATACAAATTATTAATTTATTTTTATTTTTTATTTCCCTTTGTATATATTATAAAAAAATAATCTACATAATTTTATATGATGATAATTACAATATAATATTTATAATATATATTATTTGTTAAGAGAAAAAAAAAATAAATATATACCTTCTTTTTGAGAATTGAATAAATTGTTTAATATATATATATATATATATAAATATATTATACATTTATGGTGAAAAAAAATGTTGTATTTAATTAGATTTAATATATATATATAAAAAATAGCGTATTTAAAATAATATATATATATATATATTATTATTATTACAAAATGACGGATATTATAAAAGTATATATCTATATATATGTATATATATATAATATTATTTTACTATATATATATAATATATAAATGTAAATGCATATAGTATCTATGTATATTATATATATATATAATATTAATTACATATCTAAGTTTTCTTTTCTTTTCTTTTTTTTTTTTTTTTTTTTTATTTTTTATAGAGAGCCCGTTATATATTATATATTAAAATTTTTTATAATAGCATTTATATACAATATTTTATTAACTAAAAGAAAAAAAAAAAAAAAAAAAAAAAAAAAACGAGGAAATTTATATTTCTTTAACAACATTTTAATTAAAATCATGATAATACAATTTCAAATCATTTTGTAATTATATAAATAAATATATATATATATATATATATATATATGTACTTTTAAATTAGGAATATTCTCATTTATAAATATATCTTATTTTTTAAATTGGTATAAAAAAAAAAAAAAAAATAAGAAACCGTTGATTAAATAATACATATATAATATAAATATATTTTATAAATATATATTTATATATATATATATATATATTTATAACGTATATCATTTTAAAGATAAACTAGTatggcaccaaaaaaaaaaagaaaagttacgcgtgatccaacaagaagtgcaaatagtggagcaggagcaggagcaggagcaatattaagtagagctagcatggcttctagaatcctctggcatgagatGtggcatgaaggcctggaagaggcatctcgtttgtactttggggaaaggaacgtgaaaggCatgtttgaggtgctggagcccttgcatgctatgatggaacggggcccccagactctgaaGgaaacatcctttaatcaggcctatggtcgagatttaatggaggcccaagagtggtgcagGaagtacatgaaatcagggaatgtcaaggacctcctccaagcctgggacctctattatcaTgtgttccgacgaatctcaaagGGTGAGGGTCGTGGTTCACTTCTTACTTGCGGTGACGTTGAGGAGAACCCTGGTCCTGGTACCggatcccatgcatggttcgctaaactgcatcgtcgctgtgtcccagaacatgggcatcggcaagaacggggactacccctggccaccgctcaggaacgaatttagatatttccagagaatgaccacaacctcttcagtagaaggtaaacagaatctggtgattatgggtaagaagacctggttctccattcctgagaagaatcgacctttaaagggtagaattaatttagttctcagcagagaactcaaggaacctccacaaggagctcattttctttccagaagtctagatgatgccttaaaacttactgaacaaccagaattagcaaataaagtagacatggtctggatagttggtggcagttctgtttataaggaagccatgaatcacccaggccatcttaaactatttgtgacaaggatcatgcaagactttgaaagtgacacgttttttccagaaattgatttggagaaatataaacttctgccagaatacccaggtgttctctctgatgtccaggaggagaaaggcattaagtacaaatttgaagtatatgagaagaatgattaagcttatttaataatagattaaaaatattataaaaataaaaacataaacacagaaattacaaaaaaaatacatatgaattttttttttgtaatcttccttataaatatagaataatgaatcatataaaacatatcattattcatttatttacatttaaaattattgtttcagtatctttaatttattatgtatatataaaaataacttacaattttattaataaacaatatatgtttattaattcatgttttgtaatttatgggatagcgattttttttactgtctgtatttttcttttttaattatgttttaattgtattttatttttattattgttctttttatagtattattttaaaacaaaatgtattttctaagaacttataataataataaatataaattttaataaaaattatatttatcttttacaatatgaacataaagtacaacattaatatatagcttttaatatttttattcctaatcatgtaaatcttaaatttttctttttaaacatatgttaaatatttatttctcattatatataagaacatatttattaaatctagaattctatagtgagtcgtattacaattcactggccgtcgttttacaacgtcgtgactgggaaaaccctggcgttacccaacttaatcgccttgcagcacatccccctttcgccagctggcgtaatagcgaagaggcccgcaccgatcgcccttcccaacagttgcgcagcctgaatggcgaatggcgcctgatgcggtattttctccttacgcatctgtgcggtatttcacaccgcatatggtgcactctcagtacaatctgctctgatgccgcatagttaagccagccccgacacccgccaacacccgctgacgcgccctgacgggcttgtctgctcccggcatccgcttacagacaagctgtgaccgtctccgggagctgcatgtgtcagaggttttcaccgtcatcaccgaaacgcgcgagacgaaagggcctcgtgatacgcctatttttataggttaatgtcatgataataatggtttcttagacgtcaggtggcacttttcggggaaatgtgcgcggaacccctatttgtttatttttctaaatacattcaaatatgtatccgctcatgagacaataaccctgataaatgcttcaataatattgaaaaaggaagagtatgagtattcaacatttccgtgtcgcccttattcccttttttgcggcattttgccttcctgtttttgctcacccagaaacgctggtgaaagtaaaagatgctgaagatcagttgggtgcacgagtgggttacatcgaactggatctcaacagcggtaagatccttgagagttttcgccccgaagaacgttttccaatgatgagcacttttaaagttctgctatgtggcgcggtattatcccgtattgacgccgggcaagagcaactcggtcgccgcatacactattctcagaatgacttggttgagtactcaccagtcacagaaaagcatcttacggatggcatgacagtaagagaattatgcagtgctgccataaccatgagtgataacactgcggccaacttacttctgacaacgatcggaggaccgaaggagctaaccgcttttttgcacaacatgggggatcatgtaactcgccttgatcgttgggaaccggagctgaatgaagccataccaaacgacgagcgtgacaccacgatgcctgtagcaatgccaacaacgttgcgcaaactattaactggcgaactacttactctagcttcccggcaacaattaatagactggatggaggcggataaagttgcaggaccacttctgcgctcggcccttccggctggctggtttattgctgataaatctggagccggtgagcgtgggtctcgcggtatcattgcagcactggggccagatggtaagccctcccgtatcgtagttatctacacgacggggagtcaggcaactatggatgaacgaaatagacagatcgctgagataggtgcctcactgattaagcattggtaactgtcagaccaagtttactcatatatactttagattgatttaaaacttcatttttaatttaaaaggatctaggtgaagatcctttttgataatctcatgaccaaaatcccttaacgtgagttttcgttccactgagcgtcagaccccgtagaaaagatcaaaggatcttcttgagatcctttttttctgcgcgtaatctgctgcttgcaaacaaaaaaaccaccgctaccagcggtggtttgtttgccggatcaagagctaccaactctttttccgaaggtaactggcttcagcagagcgcagataccaaatactgtccttctagtgtagccgtagttaggccaccacttcaagaactctgtagcaccgcctacatacctcgctctgctaatcctgttaccagtggctgctgccagtggcgataagtcgtgtcttaccgggttggactcaagacgatagttaccggataaggcgcagcggtcgggctgaacggggggttcgtgcacacagcccagcttggagcgaacgacctacaccgaactgagatacctacagcgtgagctatgagaaagcgccacgcttcccgaagggagaaaggcggacaggtatccggtaagcggcagggtcggaacaggagagcgcacgagggagcttccagggggaaacgcctggtatctttatagtcctgtcgggtttcgccacctctgacttgagcgtcgatttttgtgatgctcgtcaggggggcggagcctatcgaaaaacgccagcaacgcggcctttttacggttcctggccttttgctggccttttgctcacatgttctttcctgcgttatcccctgattctgtggataaccgtattaccgcctttgagtgagctgataccgctcgccgcagccgaacgaccgagcgcagcgagtcagtgagcgaggaagcggaagagcgcccaatacgcaaaccgcctctccccgcgcgttggccgattcattaatgcagctggcacgacaggtttcccgactggaaagcgggcagtgagcgcaacgcaattaatgtgagttagctcactcattaggcaccccaggctttacactttatgcttccggctcgtatgttgtgtggaattgtgagcggataacaatttcacacaggaaacagctatgaccatgattacgccaagctatttaggtgacactatagaatactcgcggccgctaaATTGATCCGGTTAAAAGAGATTTGACAAATCTTTTTTTAGAAAAGCGGGAGATGGCAAAAGAAGAGATAGACCCTCGAAAAGATTGGTTAAAGAGAATATTGTTGAATAATAACACGAACGATTTAAGGATGTTTTTAAAGGCTAGCGAATTAAAAACAAAAGAAGGTGATTATTGTATGACATGCAAAAGTAACGTAAAACAGTTATTATATCTTCATACAAAAAAGAATTATTGTCATTTATGTGAAGAAATCTTTTGTGCTTATTGTGTAAAAAGTATTGATTTTATGAAAGATGAAAAAGAGAAATATATAAAAATAAGATTATGTAGAAATTGTTTTATATATATAAATGAACTAAAATATATAATTAATCCAAATTTATCTGTAGATAGAAAAGCTATAGATTTACAAAATTCATTCAATGATATATCAAATTGTTATACAAACTTATGTAGTAACGTACCTCAATTAAATGGATTAGTATTATTATGCGAAAATAATAAAGAATTTTTAGATAGCTTTAAAAGTGAAATTAAACAGTTGGAAGAAAAGATTCAGCATGATGTTGACTTTTTAAATATCATGAAAAAAAAAAATAATTTTTTATCAGACAATACATTAATACTTAATAAAATGGGAAAAAACTTATTATTATATTTGAAAATTATAAGAAATAAAATTATTCCATCATCTATTGAAGTTCTAAATAAAACAAAGGAATTGCTATATAAGAAAcctaggTCAGGATTGAGATCAAGATCTGCTGCTGCTGGTGCTGGTGGTGCTGCTAGAGCTGCTctgcagTAGTATTCGACGTTGAATTGTTAAAGTTAGAGACATGGAAAGAAATTTGATTCATCTCGTGATAGAAATAAACCATTTAAATTTATGCTAGGTAAACAAGAAGTAATACGAGGTTGGGAAGAAGGAGTTGCTCAAATGAGTGTAGGTCAAAGAGCAAAACTTACTATATCTCCAGATTATGCTTATGGTGCAACTGGACATCCAGGTATAATTCCACCTCATGCAACTCTTGTATTTGATGTGGAGCTTCTAAAACTAGAAACTAGAGGTGTTCAGGTTGAAACAATTTCACCTGGAGATGGCAGAACCTTTCCTAAAAGAGGACAGACTTGCGTAGTTCATTATACAGGCATGCTAGAGGATGGTAAGAAATTTGATTCTAGTCGAGATAGAAATAAGCCATTCAAGTTTATGCTAGGTAAACAGGAAGTAATAAGAGGTTGGGAAGAGGGTGTAGCACAGATGTCAGTTGGACAAAGAGCAAAGTTAACAATATCACCAGATTATGCATACGGTGCAACAGGCCATCCTGGCATCATCCCTCCACATGCAACTTTAGTATTCGACGTTGAATTGTTAAAGTTAGAGACAacgcgtGCTAGAGGTGCTGCTGCTGGTGCTGGAGGTGCAGGTAGACGTACGATGAGTAAAGGAGAAGAACTTTTCACTGGAGTTGTCCCAATTCTTGTTGAATTAGATGGTGATGTTAATGGGCACAAATTTTCTGTCAGTGGAGAGGGTGAAGGTGATGCAACATACGGAAAACTTACCCTTAAATTTATTTGCACTACTGGAAAACTACCTGTTCCATGGCCAACACTTGTCACTACTTTCGCGTATGGTCTTCAATGCTTTGCGAGATACCCAGATCATATGAAACAGCATGACTTTTTCAAGAGTGCCATGCCCGAAGGTTATGTACAGGAAAGAACTATATTTTTCAAAGATGACGGGAACTACAAGACACGTGCTGAAGTCAAGTTTGAAGGTGATACCCTTGTTAATAGAATCGAGTTAAAAGGTATTGATTTTAAAGAAGATGGAAACATTCTTGGACACAAATTGGAATACAACTATAACTCACACAATGTATACATCATGGCAGACAAACAAAAGAATGGAATCAAAGTTAACTTCAAAATTAGACACAACATTGAAGATGGAAGCGTTCAACTAGCAGACCATTATCAACAAAATACTCCAATTGGCGATGGCCCTGTCCTTTTACCAGACAACCATTACCTGTCCACACAATCTGCCCTTTCGAAAGATCCCAACGAAAAGAGAGACCACATGGTCCTTCTTGAGTTTGTAACAGCTGCTGGGATTACACATGGCATGGATGAGCTCTACAAAGTCGACGGAGAAGGAAGAGGAAGTTTATTAACATGTGGAGATGTAGAAGAAAATCCAGGACCAATGATTGAACAAGATGGATTGCACGCAGGTTCTCCGGCCGCTTGGGTGGAGAGGCTATTCGGCTATGACTGGGCACAACAGACAATCGGCTGCTCTGATGCCGCCGTGTTCCGGCTGTCAGCGCAGGGGCGCCCGGTTCTTTTTGTCAAGACCGACCTGTCCGGTGCCCTGAATGAACTGCAGGACGAGGCAGCGCGGCTATCGTGGCTGGCCACGACGGGCGTTCCTTGCGCAGCTGTGCTCGACGTTGTCACTGAAGCGGGAAGGGACTGGCTGCTATTGGGCGAAGTGCCGGGGCAGGATCTCCTGTCATCTCACCTTGCTCCTGCCGAGAAAGTATCCATCATGGCTGATGCAATGCGGCGGCTGCATACGCTTGATCCGGCTACCTGCCCATTCGACCACCAAGCGAAACATCGCATCGAGCGAGCACGTACTCGGATGGAAGCCGGTCTTGTCGATCAGGATGATCTGGACGAAGAGCATCAGGGGCTCGCGCCAGCCGAACTGTTCGCCAGGCTCAAGGCGCGCATGCCCGACGGCGAGGATCTCGTCGTGACCCATGGCGATGCCTGCTTGCCGAATATCATGGTGGAAAATGGCCGCTTTTCTGGATTCATCGACTGTGGCCGGCTGGGTGTGGCGGACCGCTATCAGGACATAGCGTTGGCTACCCGTGATATTGCTGAAGAGCTTGGCGGCGAATGGGCTGACCGCTTCCTCGTGCTTTACGGTATCGCCGCTCCCGATTCGCAGCGCATCGCCTTCTATCGCCTTCTTGACGAGTTCTTCTAACTCGAG

**>SLI2a-2x-FKBP**

XXXX = INSERT (homology region of POI)

XXXX = Linker

XXXX = FKBP

XXXX = mScarlet

XXXX = T2A skip peptide

XXXX = yDHODH resistance

XXXX = BSD

atttaataatagattaaaaatattataaaaataaaaacataaacacagaaattacaaaaaaaatacatatgaattttttttttgtaatcttccttataaatatagaataatgaatcatataaaacatatcattattcatttatttacatttaaaattattgtttcagtatctttaatttattatgtatatataaaaataacttacaattttattaataaacaatatatgtttattaattcatgttttgtaatttatgggatagcgattttttttactgtctgtatttttcttttttaattatgttttaattgtattttatttttattattgttctttttatagtattattttaaaacaaaatgtattttctaagaacttataataataataaatataaattttaataaaaattatatttatcttttacaatatgaacataaagtacaacattaatatatagcttttaatatttttattcctaatcatgtaaatcttaaatttttctttttaaacatatgttaaatatttatttctcattatatataagaacatatttattaaatctagaattctatagtgagtcgtattacaattcactggccgtcgttttacaacgtcgtgactgggaaaaccctggcgttacccaacttaatcgccttgcagcacatccccctttcgccagctggcgtaatagcgaagaggcccgcaccgatcgcccttcccaacagttgcgcagcctgaatggcgaatggcgcctgatgcggtattttctccttacgcatctgtgcggtatttcacaccgcatatggtgcactctcagtacaatctgctctgatgccgcatagttaagccagccccgacacccgccaacacccgctgacgcgccctgacgggcttgtctgctcccggcatccgcttacagacaagctgtgaccgtctccgggagctgcatgtgtcagaggttttcaccgtcatcaccgaaacgcgcgagacgaaagggcctcgtgatacgcctatttttataggttaatgtcatgataataatggtttcttagacgtcaggtggcacttttcggggaaatgtgcgcggaacccctatttgtttatttttctaaatacattcaaatatgtatccgctcatgagacaataaccctgataaatgcttcaataatattgaaaaaggaagagtatgagtattcaacatttccgtgtcgcccttattcccttttttgcggcattttgccttcctgtttttgctcacccagaaacgctggtgaaagtaaaagatgctgaagatcagttgggtgcacgagtgggttacatcgaactggatctcaacagcggtaagatccttgagagttttcgccccgaagaacgttttccaatgatgagcacttttaaagttctgctatgtggcgcggtattatcccgtattgacgccgggcaagagcaactcggtcgccgcatacactattctcagaatgacttggttgagtactcaccagtcacagaaaagcatcttacggatggcatgacagtaagagaattatgcagtgctgccataaccatgagtgataacactgcggccaacttacttctgacaacgatcggaggaccgaaggagctaaccgcttttttgcacaacatgggggatcatgtaactcgccttgatcgttgggaaccggagctgaatgaagccataccaaacgacgagcgtgacaccacgatgcctgtagcaatgccaacaacgttgcgcaaactattaactggcgaactacttactctagcttcccggcaacaattaatagactggatggaggcggataaagttgcaggaccacttctgcgctcggcccttccggctggctggtttattgctgataaatctggagccggtgagcgtgggtctcgcggtatcattgcagcactggggccagatggtaagccctcccgtatcgtagttatctacacgacggggagtcaggcaactatggatgaacgaaatagacagatcgctgagataggtgcctcactgattaagcattggtaactgtcagaccaagtttactcatatatactttagattgatttaaaacttcatttttaatttaaaaggatctaggtgaagatcctttttgataatctcatgaccaaaatcccttaacgtgagttttcgttccactgagcgtcagaccccgtagaaaagatcaaaggatcttcttgagatcctttttttctgcgcgtaatctgctgcttgcaaacaaaaaaaccaccgctaccagcggtggtttgtttgccggatcaagagctaccaactctttttccgaaggtaactggcttcagcagagcgcagataccaaatactgtccttctagtgtagccgtagttaggccaccacttcaagaactctgtagcaccgcctacatacctcgctctgctaatcctgttaccagtggctgctgccagtggcgataagtcgtgtcttaccgggttggactcaagacgatagttaccggataaggcgcagcggtcgggctgaacggggggttcgtgcacacagcccagcttggagcgaacgacctacaccgaactgagatacctacagcgtgagctatgagaaagcgccacgcttcccgaagggagaaaggcggacaggtatccggtaagcggcagggtcggaacaggagagcgcacgagggagcttccagggggaaacgcctggtatctttatagtcctgtcgggtttcgccacctctgacttgagcgtcgatttttgtgatgctcgtcaggggggcggagcctatcgaaaaacgccagcaacgcggcctttttacggttcctggccttttgctggccttttgctcacatgttctttcctgcgttatcccctgattctgtggataaccgtattaccgcctttgagtgagctgataccgctcgccgcagccgaacgaccgagcgcagcgagtcagtgagcgaggaagcggaagagcgcccaatacgcaaaccgcctctccccgcgcgttggccgattcattaatgcagctggcacgacaggtttcccgactggaaagcgggcagtgagcgcaacgcaattaatgtgagttagctcactcattaggcaccccaggctttacactttatgcttccggctcgtatgttgtgtggaattgtgagcggataacaatttcacacaggaaacagctatgaccatgattacgccaagctatttaggtgacactatagaatactcgcggccgctaa**INSERT**CCTAGGGCCAGGGGAGCAGCCGCAGGAGCAGGGGGGGCAGGAAGGCGTGGTGTTCAGGTCGAGACTATTAGCCCTGGAGATGGACGCACGTTTCCTAAGCGTGGACAGACATGCGTAGTTCACTACACAGGTATGTTGGAGGACGGTAAAAAGTTCGACAGCTCACGCGACCGCAATAAACCTTTCAAGTTTATGCTTGGCAAGCAGGAGGTTATTCGTGGATGGGAGGAGGGTGTAGCACAGATGTCTGTTGGACAGCGTGCTAAGTTGACAATTTCACCTGACTATGCTTATGGCGCTACGGGCCATCCCGGGATCATTCCGCCACATGCGACTCTGGTATTCGACGTTGAATTATTAAAGTTAGAGACAgctagaggggccgctgcaggtgctggtggagctggaagaCGTGGAGTACAAGTAGAGACTATCTCTCCAGGTGACGGTCGCACTTTCCCAAAGCGTGGCCAAACCTGTGTTGTACATTACACTGGTATGCTGGAGGATGGGAAAAAGTTCGATTCCAGTCGCGACCGTAACAAACCGTTCAAATTCATGTTGGGAAAGCAGGAAGTGATCCGCGGGTGGGAGGAAGGCGTGGCGCAAATGAGCGTCGGTCAGCGGGCTAAATTGACCATTTCCCCTGACTACGCGTATGGGGCTACTGGGCACCCAGGGATTATTCCGCCTCACGCTACACTTGTGTTTGATGTCGAACTTTTGAAACTGGAAACTATGGTGAGTAAGGGTGAGGCAGTGATTAAGGAGTTTATGCGTTTTAAGGTGCACATGGAGGGTAGTATGAACGGTCACGAGTTTGAGATTGAGGGTGAGGGTGAGGGTCGTCCATACGAGGGTACACAGACAGCAAAGCTTAAGGTGACAAAGGGTGGTCCACTTCCATTTAGTTGGGATATTCTTAGTCCACAGTTTATGTACGGTAGTCGTGCATTTACAAAGCACCCAGCAGATATTCCAGATTACTACAAGCAGAGTTTTCCAGAGGGTTTTAAGTGGGAGCGTGTGATGAACTTTGAGGATGGTGGTGCAGTGACAGTGACACAGGATACAAGTCTTGAGGATGGTACACTTATTTACAAGGTGAAGCTTCGTGGTACAAACTTTCCACCAGATGGTCCAGTGATGCAGAAGAAGACAATGGGTTGGGAGGCAAGTACAGAGCGTCTTTACCCAGAGGATGGTGTGCTTAAGGGTGATATTAAGATGGCACTTCGTCTTAAGGATGGTGGTCGTTACCTTGCAGATTTTAAGACAACATACAAGGCAAAGAAGCCAGTGCAGATGCCAGGTGCATACAACGTGGATCGTAAGCTTGATATTACAAGTCACAACGAGGATTACACAGTGGTGGAGCAGTACGAGCGTAGTGAGGGTCGTCACAGTACAGGTGGTATGGATGAGCTTTACAAGGTCGACGGAGAAGGAAGAGGAAGTTTATTAACATGTGGAGATGTAGAAGAAAATCCAGGACCAATGACAGCCAGTTTAACTACCAAGTTCTTGAACAATACCTATGAAAACCCATTTATGAATGCATCCGGTGTTCATTGCATGACTACACAAGAATTAGATGAATTAGCAAACTCTAAAGCTGGCGCATTCATTACAAAGAGTGCTACAACCTTAGAAAGAGAAGGTAACCCTGAACCACGTTACATTTCTGTCCCTCTAGGCAGTATCAACTCCATGGGTTTACCAAACGAAGGTATCGACTACTATTTGTCCTATGTATTAAACCGTCAAAAGAATTATCCTGATGCACCTGCTATTTTCTTCTCAGTTGCTGGTATGAGCATTGATGAAAATTTAAATTTGTTGAGGAAAATCCAAGATAGCGAATTCAACGGTATTACCGAGTTAAACTTGTCTTGTCCTAATGTGCCTGGGAAACCACAAGTTGCTTATGACTTTGACTTGACAAAGGAAACCTTGGAAAAGGTTTTTGCCTTTTTCAAAAAACCTCTTGGTGTCAAGTTGCCTCCTTATTTTGATTTTGCCCATTTTGATATCATGGCAAAAATATTGAACGAGTTCCCATTAGCTTATGTCAACTCTATCAATAGTATAGGAAATGGTCTTTTCATTGATGTGGAGAAGGAGAGTGTAGTAGTGAAGCCAAAGAATGGTTTCGGGGGTATTGGAGGTGAATATGTTAAGCCAACCGCGCTCGCCAATGTTCGTGCATTTTACACTCGTTTGAGACCTGAAATCAAAGTTATCGGTACAGGTGGAATTAAGTCCGGTAAGGATGCATTTGAACATCTTCTATGTGGTGCCTCTATGCTACAGATTGGTACAGAATTACAAAAAGAGGGCGTCAAGATTTTTGAACGTATCGAAAAAGAATTAAAAGACATAATGGAAGCTAAGGGTTATACATCCATAGATCAGTTCCGTGGGAAGTTGAACAGCATTTAACCCGGGTCGAGGGATATGGCAGCTTAATGTTCGTTTTTCTTATTTATATATTTATACCAATTGATTGTATTTATAACTGTAAAAATGTGTATGTTGTGTGCATATTTTTTTTTGTGCATGCACATGCATGTAAATAGCTAAAATTATGAACATTTTATTTTTTGTTCAGAAAAAAAAAACTTTACACACATAAAATGGCTAGTATGAATAGCCATATTTTATATAAATTAAATCCTATGAATTTATGACCATATTAAAAATTTAGATATTTATGGAACATAATATGTTTGAAACAATAAGACAAAATTATTATTATTATTATTATTTTTACTGTTATAATTATGTGTCTCCTTCAATGATTCATAAATAGTTGGACTTGATTTTTAAAATGTTTATAATATGATTAGCATAGTTAAATAAAAAAAGTTGAAAAATTAAAAAAAAACATATAAACACAAATGATGGTTTTTCCTTCAATTTCGATATCAATTTATAGAAACAAAATATATACTTGTATAATTTTATTTTTTTATATAAATCATTACATATATAATTATACAATATTTTTTCTAAGAGATAATTATATATTAATATATATAAAAAAAGGTGTTTTTTTTTTTTTTTTTTATTTTTATTTTTATTTTATGGTAATATTTTATTTTCCTTATTTTATAAATTATATTAGTTTATATGTGATTAATTTTATATATTATCAATTTATATATTTTTAAATGCTTACTTAATTATCTTTTTTTTTTTTTTTTTTTTTTTTTCCCCTCTTTTTATATTAATTTATTTTTGAAAAAATTGATATATATATATATATATAATATATATATATACATGTAGTAGTATTAAACAATGTATAATATATATAAATAATATATTTATATATTTCATTTCAATTTTAATTTTTTTTGGTTTTTTTTTTTTTTCTTTTTGTCATATTTAAAAAAAATTATATTCATATAAGTTATGCATTTTTTATAAACATTATTCAATATATGTATAATATAATATATATATATATATTAATGTATTATTCCAATGTGCATGATAAAAGAAAAAAATAATATTTATAAAAAAAAAGAAAAATAAAACAAAAAAAGAAAAAAAAAAAAAAAAAAAAAAAAATACAAAAATAAATAATATAATTTATAATTATATATTCTTGTCACAATAAAAATATATATATATATATATATATTTATAATATGTATATTTTAAACTAGAAAAGGAATAACTAATATTTTATTTATTATCATTCAAGATTTATATTTTATAATAATAAATACCTAATAGAAATATATCAGGATCCATGGCCAAGCCTTTGTCTCAAGAAGAATCCACCCTCATTGAAAGAGCAACGGCTACAATCAACAGCATCCCCATCTCTGAAGACTACAGCGTCGCCAGCGCAGCTCTCTCTAGCGACGGCCGCATCTTCACTGGTGTCAATGTATATCATTTTACTGGGGGACCTTGTGCAGAACTCGTGGTGCTGGGCACTGCTGCTGCTGCGGCAGCTGGCAACCTGACTTGTATCGTCGCGATCGGAAATGAGAACAGGGGCATCTTGAGCCCCTGCGGACGGTGCCGACAGGTGCTTCTCGATCTGCATCCTGGGATCAAAGCGATAGTGAAGGACAGTGATGGACAGCCGACGGCAGTTGGGATTCGTGAATTGCTGCCCTCTGGTTATGTGTGGGAGGGCAAGCTT

**Insert:**

**>KIC7 homology region**

GATAGAACAAGAGATAGAACACGAGAAAAAAGTTATAAAGATAATAAAGATAGTAAATATAAAAAAAAAAAAAAATATGATGATTCAGATGATGATTTTGATGACGAATTTAGTTTAAATGATTATGATCCAAATACTCCTGGTATGATTGGAATTATGAATTCAAATAATATGAGTAATTCAGATTACAAAAATAATTTCGATAATAATTTTAATGAATTCGATGGGAATCATTTAAATTATAATCAAATGAGCGAACATAATTATCATGATTTTAATGCATTTAATAATAATATAAATAATCATATACCATTTGATAATAATAACAATCCATATTCAAAAAATAATTTTCATTTAAAATATATGAACTCACCAAGTAGTGATCTAGCTAATATCAATCCAATATGTTCAGATGATTTTAATCCAATGTATTTTAATAAAATGAATTATCCTAATAATTTTAATGTGAAATTAAATGAAGATAGTGATATAAATAATAAAATGAAATATAATAATAAAATGAGTATGGTACCATATGACCCTTTTGATAATAGTAGTAATGGTGAAAGCAATAGATATAGTAATGCTATGGATAATGCAAATTATTATAACACCACAAATAATATGAATAACATGAGTAATATGAGTAATATGAGTAACTTGAATAATATGAATAATATGAGTAACATGAATAATATGAATAATATGAGTAATATGAGTAATATGAATAATAGATTATATTTATTAAATAAAAGAAGTTCCTTAAATAATAATTATTCACCAAATAATATAAAACCTTTATATAATCAACAATATTCAAATAAAAATTTGTATAATACCCTAAATCATAATCAAAATCCATTGAACCAAAAAATGTCTTTTGATAATAATATACCGAGTAATAAAAATAAAATGTATCAAAACAAAAAAAATTCAACATATTTATTAAATAATAATTTAAATAAATCAAATAGTGGTTCAGGATTATTACACTCAGGCAATAATATGAATTTATTATGTTATTCACAACAAAATAGTTTAAATGATATGAATAGATTTCATGATAATAATATGTATAATCCTGATTTTGCCCCTGTAAACGTTTCTGATAAAAACCCATTTTCTCTACACAATATGATATCCACACCTAACAACCGAAAATCATATAATGCTTTAAATAGTTTAAATAGTGATGTAAGTAATAATAGTATAAGTAGAAATTCATTTAAGGCATTGAGTTATCCTGTTTTTGATTCAAACAAAAAATTT

**Insert:**

**>Sand1 homology region**

TAGATGGATTTGAACCTCTACCTTTAAAACCAGATTATCGTAATAAAGTACATAATCTTATATCCTCTTTTAAAATCAATAATGTCTTACTCTCATTTTTAATTATAGATGATAAAATTATAGGTTTATCATTATCAAAATATACATTAAATTCTATGGATATAATAATACTTATTAATATGATCACATCTATGAAGTCATTTAAAAATGCGGAATCATGGACCCCTATATGTTTACCGATTTACAATCCAAAgtaagttcattaaaaaatggaacaatatatcaatatatatatttatatatatattgaaatatatttaatatattatcaaatatgtcatatttttatatttctgttatttatttatttatttatttatttattttttttttttttcgtagCTTGTTCTTGTATGCTTACATAAATTATATTAAGAAGAAGATATGTTGCGTTTATATTTGCTCACATGCTTCTTCTCGGGATTTTTTCCACCTTTCAAGGCATACATCCATGATTGAATCGgttagtaaaaaaaataagtattgtatacaaacacaaacatatatatatgcattcttgagcagatttagactatatatgcattttattttaaccatatatacccctataaaacaatacatcactatatatatatatatatatatgtttttttttttttatttatttatatttatttttatatatatgtgtatttccttttagACACTAATAAGCACTGGCTGCTATGATGAAATAGTAAAGGCGTCAAACAATGCACCTTTTGTCTTACCACAAATTCCGGGAATAGATATTATACACTTATGTTATTACATTCCAAACTTAAAACAATATTATAGTTCAAAAATTAGTAATGATAAAATAAAACGTATATTCAGGGTATACCAAAAATGTGATGATATAATGAAGGATTGCAAATTACCTACACAGATTTATATAGAAAGTGAATATGAGAAATTTTATTGTATTAAAACGAATCTGTATCATTTATATTTATCAGTTCCTTTCTATGTAATAATTAATGATGAAAATATAAATGAAATTTTAAAAGTTATTGCAACATATCATAAAGAGATATTTATTAATAATATAAAGCAAATAACATCT

**>*crt*’-p40-mSca_*nmd3*’-NLS-FRB-T2A-DHODH**

XXXX = *crt’* promotor

XXXX = p40

XXXX = Linker

XXXX = mScarlet

XXXX = FRB

XXXX = NLS

XXXX = T2A skip peptide

XXXX = yDHODH resistance

XXXX = *nmd3’* promotor

CGGCCGCTAACGTAACAGACTTAGGAGGAGATCTtagtagttgagtgattctatatacatatacataaataaattacataatattataaattttttttatattatattagaatgtttactatataaaaataaatattttcttatatatttttttattttttcatatgaaaaaagaaattttttatatatttttttattatattttataaataaaaaagaacttatacctattattattattatatataaaaaatatatatttttttataaattcctatttttccgattatttatttattttttttttttattgtttaaaaatatataaaaaaaattcttatatatttaatattataagtccataaatatatataatttatataatttatatatttccatactttttttattatatatcatgagataaaataaatttcatatagaaatcgtatttatttattaatttgatatatattaataaataaatattaataataatatgttatatatatatatatttatttaattattatataggaaactatatatatatatatatattatatttttttttttgaaatataataatatataatattattcctaaaaatatctatattcttatacctgaacccttttttttttttttttttttttttgactttcgattgttcattctgtttattgtataaatatataaatatatatatattgtatattttattacatatattttttttttttgaaagttaagaattttaacttaataaaaaaagtacatatatttttaataaatgtcctccattatataaattgttatataaaagattttatataatttaaatagaattcatttataaacaaatttgtttataaaaatatatttatgtatatataatataaatatatatatttatatatatatatatatatatattactatatatatttttttttttttttccttttttttactttcccaagttgtactgcttctaagcttttttaataaacatatataatttgtacaaatattttagattatatacatgatgtatatttgaatatattttctatatatttgtggttccatttttgtatattatatataatatatttatatatatattgatatgtcaatatttgtataacacatgaagtttttgtttttttttttttttttttttaatggagaatatttaataatatatgaaaaaaattttatataaatatatatatatatatatatatatatatatatatatatatatatatgtatatatatatatttatatatacatgtatgttttttaaaaagttaaataattctatagattattttcattgtcttcacatatatgacataaatattttaaaatcgacattccgatatattatatttttagactataatatccgttaataataaatacacgcagtcatattatttattatacattcatttattattttgttttttttaatttcttacataTAActcgagATGGCTGTGGCCCAGCAGCTGCGGGCCGAGAGTGACTTTGAACAGCTTCCGGATGATGTTGCCATCTCGGCCAACATTGCTGACATCGAGGAGAAGAGAGGCTTCACCAGCCACTTTGTTTTCGTCATCGAGGTGAAGACAAAAGGAGGATCCAAGTACCTCATCTACCGCCGCTACCGCCAGTTCCATGCTTTGCAGAGCAAGCTGGAGGAGCGCTTCGGGCCAGACAGCAAGAGCAGTGCCCTGGCCTGTACCCTGCCCACACTCCCAGCCAAAGTCTACGTGGGTGTGAAACAGGAGATCGCCGAGATGCGGATACCTGCCCTCAACGCCTACATGAAGAGCCTGCTCAGCCTGCCGGTCTGGGTGCTGATGGATGAGGACGTCCGGATCTTCTTTTACCAGTCGCCCTATGACTCAGAGCAGGTGCCCCAGGCACTCCGCCCTAGGacaagttatccatatgataatccagattatgcAccagttgcaacattaggtaccATGGTGAGTAAGGGTGAGGCAGTGATTAAGGAGTTTATGCGTTTTAAGGTGCACATGGAGGGTAGTATGAACGGTCACGAGTTTGAGATTGAGGGTGAGGGTGAGGGTCGTCCATACGAGGGTACACAGACAGCAAAGCTTAAGGTGACAAAGGGTGGTCCACTTCCATTTAGTTGGGATATTCTTAGTCCACAGTTTATGTACGGTAGTCGTGCATTTACAAAGCACCCAGCAGATATTCCAGATTACTACAAGCAGAGTTTTCCAGAGGGTTTTAAGTGGGAGCGTGTGATGAACTTTGAGGATGGTGGTGCAGTGACAGTGACACAGGATACAAGTCTTGAGGATGGTACACTTATTTACAAGGTGAAGCTTCGTGGTACAAACTTTCCACCAGATGGTCCAGTGATGCAGAAGAAGACAATGGGTTGGGAGGCAAGTACAGAGCGTCTTTACCCAGAGGATGGTGTGCTTAAGGGTGATATTAAGATGGCACTTCGTCTTAAGGATGGTGGTCGTTACCTTGCAGATTTTAAGACAACATACAAGGCAAAGAAGCCAGTGCAGATGCCAGGTGCATACAACGTGGATCGTAAGCTTGATATTACAAGTCACAACGAGGATTACACAGTGGTGGAGCAGTACGAGCGTAGTGAGGGTCGTCACAGTACAGGTGGTATGGATGAGCTTTACAAGTAACCCGGGTCGAGGGATATGGCAGCTTAATGTTCGTTTTTCTTATTTATATATTTATACCAATTGattgtatttataactgtaaaaatgtgtatgttgtgtgcatatttttttttgtgcatgcacatgcatgtaaatagctaaaattatgaacattttattttttgttcagaaaaaaaaaactttacacacataaaatggctagtatgaatagccatattttatataaattaaatcctatgaatttatgaccatattaaaaatttagatatttatggaacataatatgtttgaaacaataagacaaaattattattattattattatttttactgttataattatgtgtctccttcaatgattcataaatagttggacttgatttttaaaatgtttataatatgattagcatagttaaataaaaaaagttgaaaaattaaaaaaaaacatataaacacaaatgatggtttttccttcaatttcgatatcaatttatagaaacaaaatatatacttgtataattttatttttttatataaatcattacatatataattatacaatattttttctaagagataattatatattaatatatataaaaaaaggtgttttttttttttttttttatttttatttttattttatggtaatattttattttccttattttataaattatattagtttatatgtgattaattttatatattatcaatttatatatttttaaatgcttacttaattatctttttttttttttttttttttttttcccctctttttatattaatttatttttgaaaaaattgatatatatataGTTGAAATATAAATTTCAAAAAAAATGATCACAAAATATACACTTAAATATAGGTACAATAAAAAAAAAAAATAAAAATATAATTACAAGATAATATTTTTTCCTGCTATCAAATTTTTATATATTCTCCTCAAAGAAAAAATATAAATAAATGAAGTAAATTAAAAAAAAAATTTTCTTTTTCTTCTTCTTTTGTAATTCCTTATTTATACATATTTTACTATATTTTCATAAAAATAAATTGTCATATTATATAAATATATATACCAAACCATAATTATATAGCCCTCAACATATTTTTATGATGTTTTTTCTTTTTAAATGTGATACGTAATTAAATATAATAATATATATTAAATATTATATTTTGTAATATTTACTTTCATGAGGTTTAATAAATTATAAAGAGAGAATAAAAAAAAAAAAAAAAAAAATTTATCCATATACAAATTATTAATTTATTTTTATTTTTTATTTCCCTTTGTATATATTATAAAAAAATAATCTACATAATTTTATATGATGATAATTACAATATAATATTTATAATATATATTATTTGTTAAGAGAAAAAAAAAATAAATATATACCTTCTTTTTGAGAATTGAATAAATTGTTTAATATATATATATATATATATAAATATATTATACATTTATGGTGAAAAAAAATGTTGTATTTAATTAGATTTAATATATATATATAAAAAATAGCGTATTTAAAATAATATATATATATATATATTATTATTATTACAAAATGACGGATATTATAAAAGTATATATCTATATATATGTATATATATATAATATTATTTTACTATATATATATAATATATAAATGTAAATGCATATAGTATCTATGTATATTATATATATATATAATATTAATTACATATCTAAGTTTTCTTTTCTTTTCTTTTTTTTTTTTTTTTTTTTTATTTTTTATAGAGAGCCCGTTATATATTATATATTAAAATTTTTTATAATAGCATTTATATACAATATTTTATTAACTAAAAGAAAAAAAAAAAAAAAAAAAAAAAAAAAACGAGGAAATTTATATTTCTTTAACAACATTTTAATTAAAATCATGATAATACAATTTCAAATCATTTTGTAATTATATAAATAAATATATATATATATATATATATATATATGTACTTTTAAATTAGGAATATTCTCATTTATAAATATATCTTATTTTTTAAATTGGTATAAAAAAAAAAAAAAAAATAAGAAACCGTTGATTAAATAATACATATATAATATAAATATATTTTATAAATATATATTTATATATATATATATATATATTTATAACGTATATCATTTTAAAGATAAACTAGTatggcaccaaaaaaaaaaagaaaagttacgcgtgatccaacaagaagtgcaaatagtggagcaggagcaggagcaggagcaatattaagtagagctagcatggcttctagaatcctctggcatgagatGtggcatgaaggcctggaagaggcatctcgtttgtactttggggaaaggaacgtgaaaggCatgtttgaggtgctggagcccttgcatgctatgatggaacggggcccccagactctgaaGgaaacatcctttaatcaggcctatggtcgagatttaatggaggcccaagagtggtgcagGaagtacatgaaatcagggaatgtcaaggacctcctccaagcctgggacctctattatcaTgtgttccgacgaatctcaaagGGTGAGGGTCGTGGTTCACTTCTTACTTGCGGTGACGTTGAGGAGAACCCTGGTCCTGTCGACATGACAGCCAGTTTAACTACCAAGTTCTTGAACAATACCTATGAAAACCCATTTATGAATGCATCCGGTGTTCATTGCATGACTACACAAGAATTAGATGAATTAGCAAACTCTAAAGCTGGCGCATTCATTACAAAGAGTGCTACAACCTTAGAAAGAGAAGGTAACCCTGAACCACGTTACATTTCTGTCCCTCTAGGCAGTATCAACTCCATGGGTTTACCAAACGAAGGTATCGACTACTATTTGTCCTATGTATTAAACCGTCAAAAGAATTATCCTGATGCACCTGCTATTTTCTTCTCAGTTGCTGGTATGAGCATTGATGAAAATTTAAATTTGTTGAGGAAAATCCAAGATAGCGAATTCAACGGTATTACCGAGTTAAACTTGTCTTGTCCTAATGTGCCTGGGAAACCACAAGTTGCTTATGACTTTGACTTGACAAAGGAAACCTTGGAAAAGGTTTTTGCCTTTTTCAAAAAACCTCTTGGTGTCAAGTTGCCTCCTTATTTTGATTTTGCCCATTTTGATATCATGGCAAAAATATTGAACGAGTTCCCATTAGCTTATGTCAACTCTATCAATAGTATAGGAAATGGTCTTTTCATTGATGTGGAGAAGGAGAGTGTAGTAGTGAAGCCAAAGAATGGTTTCGGGGGTATTGGAGGTGAATATGTTAAGCCAACCGCGCTCGCCAATGTTCGTGCATTTTACACTCGTTTGAGACCTGAAATCAAAGTTATCGGTACAGGTGGAATTAAGTCCGGTAAGGATGCATTTGAACATCTTCTATGTGGTGCCTCTATGCTACAGATTGGTACAGAATTACAAAAAGAGGGCGTCAAGATTTTTGAACGTATCGAAAAAGAATTAAAAGACATAATGGAAGCTAAGGGTTATACATCCATAGATCAGTTCCGTGGGAAGTTGAACAGCATTTAAAAGCTTATTTAATAATAGATTAAAAATATTATAAAAATAAAAACATAAACACAGAAATTACAAAAAAAATACATATGAATTTTTTTTTTGTAATCTTCCTTATAAATATAGAATAATGAATCATATAAAACATATCATTATTCATTTATTTACATTTAAAATTATTGTTTCAGTATCTTTAATTTATTATGTATATATAAAAATAACTTACAATTTTATTAATAAACAATATATGTTTATTAATTCATGTTTTGTAATTTATGGGATAGCGATTTTTTTTACTGTCTGTATTTTTCTTTTTTAATTATGTTTTAATTGTATTTTATTTTTATTATTGTTCTTTTTATAGTATTATTTTAAAACAAAATGTATTTTCTAAGAACTTATAATAATAATAAATATAAATTTTAATAAAAATTATATTTATCTTTTACAATATGAACATAAAGTACAACATTAATATATAGCTTTTAATATTTTTATTCCTAATCATGTAAATCTTAAATTTTTCTTTTTAAACATATGTTAAATATTTATTTCTCATTATATATAAGAACATATTTATTAAATCTAGAATTCTATAGTGAGTCGTATTACAATTCACTGGCCGTCGTTTTACAACGTCGTGACTGGGAAAACCCTGGCGTTACCCAACTTAATCGCCTTGCAGCACATCCCCCTTTCGCCAGCTGGCGTAATAGCGAAGAGGCCCGCACCGATCGCCCTTCCCAACAGTTGCGCAGCCTGAATGGCGAATGGCGCCTGATGCGGTATTTTCTCCTTACGCATCTGTGCGGTATTTCACACCGCATATGGTGCACTCTCAGTACAATCTGCTCTGATGCCGCATAGTTAAGCCAGCCCCGACACCCGCCAACACCCGCTGACGCGCCCTGACGGGCTTGTCTGCTCCCGGCATCCGCTTACAGACAAGCTGTGACCGTCTCCGGGAGCTGCATGTGTCAGAGGTTTTCACCGTCATCACCGAAACGCGCGAGACGAAAGGGCCTCGTGATACGCCTATTTTTATAGGTTAATGTCATGATAATAATGGTTTCTTAGACGTCAGGTGGCACTTTTCGGGGAAATGTGCGCGGAACCCCTATTTGTTTATTTTTCTAAATACATTCAAATATGTATCCGCTCATGAGACAATAACCCTGATAAATGCTTCAATAATATTGAAAAAGGAAGAGTATGAGTATTCAACATTTCCGTGTCGCCCTTATTCCCTTTTTTGCGGCATTTTGCCTTCCTGTTTTTGCTCACCCAGAAACGCTGGTGAAAGTAAAAGATGCTGAAGATCAGTTGGGTGCACGAGTGGGTTACATCGAACTGGATCTCAACAGCGGTAAGATCCTTGAGAGTTTTCGCCCCGAAGAACGTTTTCCAATGATGAGCACTTTTAAAGTTCTGCTATGTGGCGCGGTATTATCCCGTATTGACGCCGGGCAAGAGCAACTCGGTCGCCGCATACACTATTCTCAGAATGACTTGGTTGAGTACTCACCAGTCACAGAAAAGCATCTTACGGATGGCATGACAGTAAGAGAATTATGCAGTGCTGCCATAACCATGAGTGATAACACTGCGGCCAACTTACTTCTGACAACGATCGGAGGACCGAAGGAGCTAACCGCTTTTTTGCACAACATGGGGGATCATGTAACTCGCCTTGATCGTTGGGAACCGGAGCTGAATGAAGCCATACCAAACGACGAGCGTGACACCACGATGCCTGTAGCAATGCCAACAACGTTGCGCAAACTATTAACTGGCGAACTACTTACTCTAGCTTCCCGGCAACAATTAATAGACTGGATGGAGGCGGATAAAGTTGCAGGACCACTTCTGCGCTCGGCCCTTCCGGCTGGCTGGTTTATTGCTGATAAATCTGGAGCCGGTGAGCGTGGGTCTCGCGGTATCATTGCAGCACTGGGGCCAGATGGTAAGCCCTCCCGTATCGTAGTTATCTACACGACGGGGAGTCAGGCAACTATGGATGAACGAAATAGACAGATCGCTGAGATAGGTGCCTCACTGATTAAGCATTGGTAACTGTCAGACCAAGTTTACTCATATATACTTTAGATTGATTTAAAACTTCATTTTTAATTTAAAAGGATCTAGGTGAAGATCCTTTTTGATAATCTCATGACCAAAATCCCTTAACGTGAGTTTTCGTTCCACTGAGCGTCAGACCCCGTAGAAAAGATCAAAGGATCTTCTTGAGATCCTTTTTTTCTGCGCGTAATCTGCTGCTTGCAAACAAAAAAACCACCGCTACCAGCGGTGGTTTGTTTGCCGGATCAAGAGCTACCAACTCTTTTTCCGAAGGTAACTGGCTTCAGCAGAGCGCAGATACCAAATACTGTCCTTCTAGTGTAGCCGTAGTTAGGCCACCACTTCAAGAACTCTGTAGCACCGCCTACATACCTCGCTCTGCTAATCCTGTTACCAGTGGCTGCTGCCAGTGGCGATAAGTCGTGTCTTACCGGGTTGGACTCAAGACGATAGTTACCGGATAAGGCGCAGCGGTCGGGCTGAACGGGGGGTTCGTGCACACAGCCCAGCTTGGAGCGAACGACCTACACCGAACTGAGATACCTACAGCGTGAGCTATGAGAAAGCGCCACGCTTCCCGAAGGGAGAAAGGCGGACAGGTATCCGGTAAGCGGCAGGGTCGGAACAGGAGAGCGCACGAGGGAGCTTCCAGGGGGAAACGCCTGGTATCTTTATAGTCCTGTCGGGTTTCGCCACCTCTGACTTGAGCGTCGATTTTTGTGATGCTCGTCAGGGGGGCGGAGCCTATCGAAAAACGCCAGCAACGCGGCCTTTTTACGGTTCCTGGCCTTTTGCTGGCCTTTTGCTCACATGTTCTTTCCTGCGTTATCCCCTGATTCTGTGGATAACCGTATTACCGCCTTTGAGTGAGCTGATACCGCTCGCCGCAGCCGAACGACCGAGCGCAGCGAGTCAGTGAGCGAGGAAGCGGAAGAGCGCCCAATACGCAAACCGCCTCTCCCCGCGCGTTGGCCGATTCATTAATGCAGCTGGCACGACAGGTTTCCCGACTGGAAAGCGGGCAGTGAGCGCAACGCAATTAATGTGAGTTAGCTCACTCATTAGGCACCCCAGGCTTTACACTTTATGCTTCCGGCTCGTATGTTGTGTGGAATTGTGAGCGGATAACAATTTCACACAGGAAACAGCTATGACCATGATTACGCCAAGCTATTTAGGTGACACTATAGAATACTC

**>*sf3a2*’-POI-yDHODH**

XXXX = *sf3a2’* promotor

XXXX = Insert (POI)

XXXX = Linker

XXXX = mCherry

XXXX = yDHODH resistance

CGGCCGCTAACGTAACAGACTTAGGAGGAGATCTtaaAGTCTCCTTTTCTTTATTTTACAGTTGTGAATTATATTCATCTCATTTATTCTTTCTTTTTTAATAATTCAAGGTAAACGTTTTTATATTTTATCATATTATTATGGTTCATAACGTGTTGTATATATATATATATATATATATTATATATATTTATTTTTTTTTTAATAATAAAATTTTAAATTATAAAAATAAAAAAAACAAAAAAATATAATAATTTAATAAAATATAAACCTAAATATGTAATTAATTATTCTCTTTATTTTAAATTAAAAAGGAAAAAATATATATAATAAATAATGGTAAGGAAGAATCCTTATAAAATAAATATACATATAATTTTATTGAGTTAATATATAATAAGTATTATAAAAATTATATATATGTATATATATAAATATATGTTATATATATATATTTATTATATATACATAATATTATAGAATATTGACGGAAAAATATTAAAAAAAAAAAAAAAAAAAATTATATAAAATATAATTCTAAAAATTTCAGGATCCCTATTTATAAAAAAAACAAATTAACTTCAAATTCTTTTTCTTTTTTTTCATGTATATATATATATATATATATATATATATATATAATTTTTCAAAATTTTTTTAAGATATAATAAATATATTATGTATATTTTATCTATAACATTTATATTTTAAATTGAAAAAAAATAAAATAATACATAAAAGATAAATATAATGAAGTAACAATATTATTATATAATTAAATAATTGAATGAATATATATATATATATATATATATGTTAAAATTGATGAATGAATTATAATGAAAGTTAACTAAATGTAAAATATAAGAATAATATATATATTATCGAAGAAGCAATAAAATTTTTATGATATTTAGATTTCAAGTAAAACATGTATGAATAATATACATATATATATATATATATATATATATATATATATATATATATATGTACATATATACATATACATAATATATAAAATATAATTTTAAAAGTAActcgag**INSERT**CCTAGGacaagttatccatatgataatccagattatgcAccagttgcaacattaggtaccatggtgagcaagggcgaggaggataacatggccatcatCaaggagttcatgcgcttcaaggtgcacatggagggctccgtgaacggccacgagttcgaGatcgagggcgagggcgagggccgcccctacgagggcacccagaccgccaagctgaaggtGaccaagggtggccccctgcccttcgcctgggacatcctgtcccctcagttcatgtacggCtccaaggcctacgtgaagcaccccgccgacatccccgactacttgaagctgtccttcccCgagggcttcaagtgggagcgcgtgatgaacttcgaggacggcggcgtggtgaccgtgacCcaggactcctccctgcaggacggcgagttcatctacaaggtgaagctgcgcggcaccaaCttcccctccgacggccccgtaatgcagaagaagaccatgggctgggaggcctcctccgaGcggatgtaccccgaggacggcgccctgaagggcgagatcaagcagaggctgaagctgaaGgacggcggccactacgacgctgaggtcaagaccacctacaaggccaagaagcccgtgcaGctgcccggcgcctacaacgtcaacatcaagttggacatcacctcccacaacgaggactaCaccatcgtggaacagtacgaacgcgccgagggccgccactccaccggcggcatggacgagctgtacaagTAACCCGGGTCGAGGGATATGGCAGCTTAATGTTCGTTTTTCTTATTTATATATTTATACCAATTGATTGTATTTATAACTGTAAAAATGTGTATGTTGTGTGCATATTTTTTTTTGTGCATGCACATGCATGTAAATAGCTAAAATTATGAACATTTTATTTTTTGTTCAGAAAAAAAAAACTTTACACACATAAAATGGCTAGTATGAATAGCCATATTTTATATAAATTAAATCCTATGAATTTATGACCATATTAAAAATTTAGATATTTATGGAACATAATATGTTTGAAACAATAAGACAAAATTATTATTATTATTATTATTTTTACTGTTATAATTATGTGTCTCCTTCAATGATTCATAAATAGTTGGACTTGATTTTTAAAATGTTTATAATATGATTAGCATAGTTAAATAAAAAAAGTTGAAAAATTAAAAAAAAACATATAAACACAAATGATGGTTTTTCCTTCAATTTCGATATCAATTTATAGAAACAAAATATATACTTGTATAATTTTATTTTTTTATATAAATCATTACATATATAATTATACAATATTTTTTCTAAGAGATAATTATATATTAATATATATAAAAAAAGGTGTTTTTTTTTTTTTTTTTTATTTTTATTTTTATTTTATGGTAATATTTTATTTTCCTTATTTTATAAATTATATTAGTTTATATGTGATTAATTTTATATATTATCAATTTATATATTTTTAAATGCTTACTTAATTATCTTTTTTTTTTTTTTTTTTTTTTTTTCCCCTCTTTTTATATTAATTTATTTTTGAAAAAATTGATATATATATATATATATAATATATATATATACATGTAGTAGTATTAAACAATGTATAATATATATAAATAATATATTTATATATTTCATTTCAATTTTAATTTTTTTTGGTTTTTTTTTTTTTTCTTTTTGTCATATTTAAAAAAAATTATATTCATATAAGTTATGCATTTTTTATAAACATTATTCAATATATGTATAATATAATATATATATATATATTAATGTATTATTCCAATGTGCATGATAAAAGAAAAAAATAATATTTATAAAAAAAAAGAAAAATAAAACAAAAAAAGAAAAAAAAAAAAAAAAAAAAAAAAATACAAAAATAAATAATATAATTTATAATTATATATTCTTGTCACAATAAAAATATATATATATATATATATATTTATAATATGTATATTTTAAACTAGAAAAGGAATAACTAATATTTTATTTATTATCATTCAAGATTTATATTTTATAATAATAAATACCTAATAGAAATATATCAGGATCCATGACAGCCAGTTTAACTACCAAGTTCTTGAACAATACCTATGAAAACCCATTTATGAATGCATCCGGTGTTCATTGCATGACTACACAAGAATTAGATGAATTAGCAAACTCTAAAGCTGGCGCATTCATTACAAAGAGTGCTACAACCTTAGAAAGAGAAGGTAACCCTGAACCACGTTACATTTCTGTCCCTCTAGGCAGTATCAACTCCATGGGTTTACCAAACGAAGGTATCGACTACTATTTGTCCTATGTATTAAACCGTCAAAAGAATTATCCTGATGCACCTGCTATTTTCTTCTCAGTTGCTGGTATGAGCATTGATGAAAATTTAAATTTGTTGAGGAAAATCCAAGATAGCGAATTCAACGGTATTACCGAGTTAAACTTGTCTTGTCCTAATGTGCCTGGGAAACCACAAGTTGCTTATGACTTTGACTTGACAAAGGAAACCTTGGAAAAGGTTTTTGCCTTTTTCAAAAAACCTCTTGGTGTCAAGTTGCCTCCTTATTTTGATTTTGCCCATTTTGATATCATGGCAAAAATATTGAACGAGTTCCCATTAGCTTATGTCAACTCTATCAATAGTATAGGAAATGGTCTTTTCATTGATGTGGAGAAGGAGAGTGTAGTAGTGAAGCCAAAGAATGGTTTCGGGGGTATTGGAGGTGAATATGTTAAGCCAACCGCGCTCGCCAATGTTCGTGCATTTTACACTCGTTTGAGACCTGAAATCAAAGTTATCGGTACAGGTGGAATTAAGTCCGGTAAGGATGCATTTGAACATCTTCTATGTGGTGCCTCTATGCTACAGATTGGTACAGAATTACAAAAAGAGGGCGTCAAGATTTTTGAACGTATCGAAAAAGAATTAAAAGACATAATGGAAGCTAAGGGTTATACATCCATAGATCAGTTCCGTGGGAAGTTGAACAGCATTTAAAAGCTTATTTAATAATAGATTAAAAATATTATAAAAATAAAAACATAAACACAGAAATTACAAAAAAAATACATATGAATTTTTTTTTTGTAATCTTCCTTATAAATATAGAATAATGAATCATATAAAACATATCATTATTCATTTATTTACATTTAAAATTATTGTTTCAGTATCTTTAATTTATTATGTATATATAAAAATAACTTACAATTTTATTAATAAACAATATATGTTTATTAATTCATGTTTTGTAATTTATGGGATAGCGATTTTTTTTACTGTCTGTATTTTTCTTTTTTAATTATGTTTTAATTGTATTTTATTTTTATTATTGTTCTTTTTATAGTATTATTTTAAAACAAAATGTATTTTCTAAGAACTTATAATAATAATAAATATAAATTTTAATAAAAATTATATTTATCTTTTACAATATGAACATAAAGTACAACATTAATATATAGCTTTTAATATTTTTATTCCTAATCATGTAAATCTTAAATTTTTCTTTTTAAACATATGTTAAATATTTATTTCTCATTATATATAAGAACATATTTATTAAATCTAGAATTCTATAGTGAGTCGTATTACAATTCACTGGCCGTCGTTTTACAACGTCGTGACTGGGAAAACCCTGGCGTTACCCAACTTAATCGCCTTGCAGCACATCCCCCTTTCGCCAGCTGGCGTAATAGCGAAGAGGCCCGCACCGATCGCCCTTCCCAACAGTTGCGCAGCCTGAATGGCGAATGGCGCCTGATGCGGTATTTTCTCCTTACGCATCTGTGCGGTATTTCACACCGCATATGGTGCACTCTCAGTACAATCTGCTCTGATGCCGCATAGTTAAGCCAGCCCCGACACCCGCCAACACCCGCTGACGCGCCCTGACGGGCTTGTCTGCTCCCGGCATCCGCTTACAGACAAGCTGTGACCGTCTCCGGGAGCTGCATGTGTCAGAGGTTTTCACCGTCATCACCGAAACGCGCGAGACGAAAGGGCCTCGTGATACGCCTATTTTTATAGGTTAATGTCATGATAATAATGGTTTCTTAGACGTCAGGTGGCACTTTTCGGGGAAATGTGCGCGGAACCCCTATTTGTTTATTTTTCTAAATACATTCAAATATGTATCCGCTCATGAGACAATAACCCTGATAAATGCTTCAATAATATTGAAAAAGGAAGAGTATGAGTATTCAACATTTCCGTGTCGCCCTTATTCCCTTTTTTGCGGCATTTTGCCTTCCTGTTTTTGCTCACCCAGAAACGCTGGTGAAAGTAAAAGATGCTGAAGATCAGTTGGGTGCACGAGTGGGTTACATCGAACTGGATCTCAACAGCGGTAAGATCCTTGAGAGTTTTCGCCCCGAAGAACGTTTTCCAATGATGAGCACTTTTAAAGTTCTGCTATGTGGCGCGGTATTATCCCGTATTGACGCCGGGCAAGAGCAACTCGGTCGCCGCATACACTATTCTCAGAATGACTTGGTTGAGTACTCACCAGTCACAGAAAAGCATCTTACGGATGGCATGACAGTAAGAGAATTATGCAGTGCTGCCATAACCATGAGTGATAACACTGCGGCCAACTTACTTCTGACAACGATCGGAGGACCGAAGGAGCTAACCGCTTTTTTGCACAACATGGGGGATCATGTAACTCGCCTTGATCGTTGGGAACCGGAGCTGAATGAAGCCATACCAAACGACGAGCGTGACACCACGATGCCTGTAGCAATGCCAACAACGTTGCGCAAACTATTAACTGGCGAACTACTTACTCTAGCTTCCCGGCAACAATTAATAGACTGGATGGAGGCGGATAAAGTTGCAGGACCACTTCTGCGCTCGGCCCTTCCGGCTGGCTGGTTTATTGCTGATAAATCTGGAGCCGGTGAGCGTGGGTCTCGCGGTATCATTGCAGCACTGGGGCCAGATGGTAAGCCCTCCCGTATCGTAGTTATCTACACGACGGGGAGTCAGGCAACTATGGATGAACGAAATAGACAGATCGCTGAGATAGGTGCCTCACTGATTAAGCATTGGTAACTGTCAGACCAAGTTTACTCATATATACTTTAGATTGATTTAAAACTTCATTTTTAATTTAAAAGGATCTAGGTGAAGATCCTTTTTGATAATCTCATGACCAAAATCCCTTAACGTGAGTTTTCGTTCCACTGAGCGTCAGACCCCGTAGAAAAGATCAAAGGATCTTCTTGAGATCCTTTTTTTCTGCGCGTAATCTGCTGCTTGCAAACAAAAAAACCACCGCTACCAGCGGTGGTTTGTTTGCCGGATCAAGAGCTACCAACTCTTTTTCCGAAGGTAACTGGCTTCAGCAGAGCGCAGATACCAAATACTGTCCTTCTAGTGTAGCCGTAGTTAGGCCACCACTTCAAGAACTCTGTAGCACCGCCTACATACCTCGCTCTGCTAATCCTGTTACCAGTGGCTGCTGCCAGTGGCGATAAGTCGTGTCTTACCGGGTTGGACTCAAGACGATAGTTACCGGATAAGGCGCAGCGGTCGGGCTGAACGGGGGGTTCGTGCACACAGCCCAGCTTGGAGCGAACGACCTACACCGAACTGAGATACCTACAGCGTGAGCTATGAGAAAGCGCCACGCTTCCCGAAGGGAGAAAGGCGGACAGGTATCCGGTAAGCGGCAGGGTCGGAACAGGAGAGCGCACGAGGGAGCTTCCAGGGGGAAACGCCTGGTATCTTTATAGTCCTGTCGGGTTTCGCCACCTCTGACTTGAGCGTCGATTTTTGTGATGCTCGTCAGGGGGGCGGAGCCTATCGAAAAACGCCAGCAACGCGGCCTTTTTACGGTTCCTGGCCTTTTGCTGGCCTTTTGCTCACATGTTCTTTCCTGCGTTATCCCCTGATTCTGTGGATAACCGTATTACCGCCTTTGAGTGAGCTGATACCGCTCGCCGCAGCCGAACGACCGAGCGCAGCGAGTCAGTGAGCGAGGAAGCGGAAGAGCGCCCAATACGCAAACCGCCTCTCCCCGCGCGTTGGCCGATTCATTAATGCAGCTGGCACGACAGGTTTCCCGACTGGAAAGCGGGCAGTGAGCGCAACGCAATTAATGTGAGTTAGCTCACTCATTAGGCACCCCAGGCTTTACACTTTATGCTTCCGGCTCGTATGTTGTGTGGAATTGTGAGCGGATAACAATTTCACACAGGAAACAGCTATGACCATGATTACGCCAAGCTATTTAGGTGACACTATAGAATACTC

**Insert:**

**>Rab5b**

ATGGGATGTTCATCAAGCACCGAAAGGTTAACATCTACCAAAAATATTAATATTGTTACATCTCCTGCACAACAACAAAAAAAAAATGCTCAAGATACTAAAGTGAAAATAGTTTTATTAGGTGACAGTGGTGTTGGAAAATCAAGTATTGCTTTGTATTTATGTCATGGTCGCTTTTCTGAAAAACATCAAGTTACTATAGGTGCAGCATTTTTACATCATAATATTGAATTAAAAAATGgtaaaaattaagagacataaaaataaattgtatgtctgaatatgaataaaatgtttttagagaatttttctatacatacacgcacatatatatatatatatatatatatatatatatttattttattattattatctttttattgtatttaagGTGCAACAATGAAATTACATATTTGGGATACTGGAGGGCAAGAAAGATTTCGTTCCATGGCTCCTTTATATTATCGTGACGCATATGGAGCAGTTGTCGTTTATGACTCAAAgtaatattttaatcattgaataattattttctatttttttttttttttaatataaccaaattggctagctgttgtcatatttatatatacataaaatttttttttttttttttttttttttaaattcgctagctgttcatatattatataaatatacatttatatatacataattattttttttttttttatgaaaaaatatcaaagTAATGTCGAATCCTTTGATTCTTTAAAATACTGGATAAACGAAATAAAATCAAATGGTCCAAGAAATTGTTGTATCATGGTAGTTGCCAATAAAAAGGACTTACCCCAAAAACTGAATTCAGAGgtaatcatgaggacacattaaaaagtttatatattcatatagataagaaaaatgatatatatatatatatatatatatatatatatatatatttatatttatatatttatatatgtatgtcaatttatttaactttttattttattagATGGTCATGAAGTTTTGTGAACAAGAAAATGTATCATTTATTGAATGCTCAGCTAAGgtataaatattcaaaaaaaatatattaataaatttataacatattataagaatagcaacatataagagttttacaattatttgattccctctctataaatatatatatatatatatatatatatatttttttagACAGGAGAGAATATTACTACTTTATTTGAAAAATTAGgttagattatttataatttttacatatttccaatgaaaataattgaataattaaataaaaaaataaaatctataatactacaataatatcttatttatattttatattatattatattatattatattatattatattatattatattatattatattatattatattatattattttttattttttgttttacagCTAGTCGCATTTATTCCCGATTTAAAGAAGTTTTATATTATAACAATCCT

**Insert:**

**>VPS45**

ATGGAGAATAATCCTTACGTGTTTAAAAGCTTAGTTAACATTTATGAGGAATATTTAAATTTAATTATTCATCGTGTGAAGGGATATAAAGTATTAGTGCTTGATGATGAAACGAAGAGTATTATATCGTTGATTTTTTCACATTCGTATATATTGGAGAAGGAGATATTTTTAACATTAAATTTTAATGATAAGAACATATTTGAAGATATATATAATAATAATAATGATAAGAAAGAAAATTTTGATTTTATGAATTATAAGATAAAGAATTTAAAACATTTGAAAGTTATCTTTTTACTAAGACCTACCTATACGAATATATTAAGATTAATGAGTGAACTTAAGAAACCGTTATTTTCAGAATATTATATATTTTTTACAAATACAATAAATGATATATATATAGAGAAGTTAGCTAAGGCGGATGAGTTTGATGTAATAAAAAATATAATAGAATATTATATTGATACATATGTATTGCATGATTATTTATTTCATCTAAATATAGACTACACTTCCTTTTTATATAAGAATGATCATAAATTTATAGACAAAGAAAAGAAAAAAAAAGAGTTAAATTATTTTAAGCAATATAATAATAATAATAATATTAATAGTAATAATAATTATTCTAGTGATGGTCGCTATGAAAAGCTAACCATTGAAGAGTTTAACAAATTAGAAGGAAATAATAATATGATATATGATAATAATAATAATAATAATAATAATAATAATATTAATAGTGGTAATATTAATTATAGTCATTTTAATTTATCTATTGAACATATTAATAACGATAATAGGAACAATTCAAATATAACGCTTTATATGAATCAAATAGTTCAACGAATTATCGATGGATTGTTTTCTTTCCTTTGTTGTATAAGACAAGTTCCTGATGTTATTTATAACAGGCATTCTAAAATATGTAAACATATTATAGATATGTTAAAAGAAAAAATGTTAAGACATCAATCTGTATTTAATAATATATTAGATATATATGAAAAATATAATGATGAAATGGAGAGGAAAAAAAAGAAGAAGATATTAGAAACAAATAATGAACCTAATTACCAATTTAATCATTTAATAAATCAAAATATACATGAGATAACAGAAGGTGATGCTTGTTATTTTTTAATTTTAGATAGAAATGAAGATCCTATAACACCTTTACTTACACAATGGACATATCAATCTATGTTACATGAACTTATAGGAATAGAAAATAATAAAATAAATTTAAATTGTAATAATAAAGAAGAAGAACAACAACAAATTGTTATGTCTTGTAATTATGATGATTTTTATAATGAACATTTATTTGATAACTTTGGAGATTTAGGTCAGGCTGTTAAAAATTATGTAGATATTTATCAAGAGGAAACTTCAAAAAAAACAAATTTAGAATCTATTGATGATATACAAAAATTTATAGATATATATCCAAATTATAAAAAATTATCAGGAAATGTAACAAAACATGTTAATATTTTACACAAATTTTCTGATATAGTACAAAAAAGACAACTATTTTATATTTCTGAGTTAGAACAATCAATAGCTTGTTATCATACAAAAAATGATCATTTTAAACAAGTTATTGATACTATTAAAAATTATACCTATACTAATTATGATGTCTTACGATTATCTTTATTATATTCTTTAAAATATGCAGATGAACAACATATCAATGTTATAAAAAATGAACTAGCTAAAAGAAATATACAAAAAGATCAAATTTTATTAATAGATGCTTTATTATTATATTCTAGTCAACAAACAAAATATAATCAATTATTCAAAGAACAAACCTTTCTAAACCTAGCCAAAACAACTATTACAAGAACTATCAAAGGAACATCAAATGTATTTACATTACATAAATCTTATCTTTATTATTTATTAGAAGATATTATAAAATATAAAATTAATACTCAATTATATACAACCACAAACTTGTTACACACAGAACCTACTTTGAATAAAAAAATCAATTCCATTGTCGTTTTTTTTATAGGAGGTGCTACATATGAAGAATATAGAGATGTACAACATTTGAGTAAAAAATATAACATATCCATTGTCTTGGGTTCAACGCACATGCATAATTCACAGTCATTTCTTGCAGACGTTTTGCAGCTTATCAAGAAA

**>*sf3a2*’-2xFYVE(of Rbsn5L)-yDHODH**

XXXX = *sf3a2’* promotor

XXX = FYVE

XXXX = yDHODH

XXXX = mCherry

XXXXX = linker

CGGCCGCTAACGTAACAGACTTAGGAGGAGATCTttattattattacatTAAAGTCTCCTTTTCTTTATTTTACAGTTGTGAATTATATTCATCTCATTTATTCTTTCTTTTTTAATAATTCAAGGTAAACGTTTTTATATTTTATCATATTATTATGGTTCATAACGTGTTGTATATATATATATATATATTATATATATTTATTTTTTTTTTAATAATAAAATTTTAAATTATAAAAATAAAAAAAACAAAAAAATATAATAATTTAATAAAATATAAACCTAAATATGTAATTAATTATTCTCTTTATTTTAAATTAAAAAGGAAAAGATATATATAATAAATAATGGTAAGGAAGAATCCTTATAAAATAAATATACATATAATTTTATTGAGTTAATATATAATAAGTATTATAAAAATTATATATATGTATATATATAAATATATGTTATATATATATATTTATTATATATACATAATATTATAGAATATTGACGGAAAAATATTAAAAAAAAAAAAAAAATTATATAAAATATAATTCTAAAAATTTCAGGATCCCTATTTATAAAAAAAACAAATTAACTTCAAATTCTTTTTCTTTTTTTTCATGTATATATATATATATATATATATATATATAATTTTTCAAAATTTTTTTAAGATATAATAAATATATTATGTATATTTTATCTATAACATTTATATTTTAAATTGAAAAAAAATAAAATAATACATAAAAGATAAATATAATGAAGTAACAATATTATTATATAATTAAATAATTGAATGAATATATATATATATATATATATATATGTTAAAATTGATGAATGAATTATAATGAAAGTTAACTAAATGTAAAATATAAGAATAATATATATATTATCGAAGAAGCAATAAAATTTTTATGATATTTAGATTTCAAGTAAAACATGTATGAATAATATACATATATATATATATATATATATATATATATATATATATATATATGTACATATATACATATACATAATATATAAAATATAATTTTAAAAGtAActcgagatggtgagcaagggcgaggaggataacatggccatcatCaaggagttcatgcgcttcaaggtgcacatggagggctccgtgaacggccacgagttcgaGatcgagggcgagggcgagggccgcccctacgagggcacccagaccgccaagctgaaggtGaccaagggtggccccctgcccttcgcctgggacatcctgtcccctcagttcatgtacggCtccaaggcctacgtgaagcaccccgccgacatccccgactacttgaagctgtccttcccCgagggcttcaagtgggagcgcgtgatgaacttcgaggacggcggcgtggtgaccgtgacCcaggactcctccctgcaggacggcgagttcatctacaaggtgaagctgcgcggcaccaaCttcccctccgacggccccgtaatgcagaagaagaccatgggctgggaggcctcctccgaGcggatgtaccccgaggacggcgccctgaagggcgagatcaagcagaggctgaagctgaaGgacggcggccactacgacgctgaggtcaagaccacctacaaggccaagaagcccgtgcaGctgcccggcgcctacaacgtcaacatcaagttggacatcacctcccacaacgaggactaCaccatcgtggaacagtacgaacgcgccgagggccgccactccaccggcggcatggacgagctgtacaagacgcgtgatccaacaagaagtgcaaatagtggagcaggagcaggagcaggagcaatattaagtagagctagcATAGACCCTCGAAAAGATTGGTTAAAGAGAATATTGTTGAATAATAACACGAACGATTTAAGGATGTTTTTAAAGGCTAGCGAATTAAAAACAAAAGAAGGTGATTATTGTATGACATGCAAAAGTAACGTAAAACAGTTATTATATCTTCATACAAAAAAGAATTATTGTCATTTATGTGAAGAAATCTTTTGTGCTTATTGTGTAAAAAGTATTGATTTTATGAAAGATGAAAAAGAGAAATATATAAAAATAAGATTATGTAGAAATTGTTTTATATATATAAATGAACTAAAATATATAATTAATCCAacgcgtgatccaacaagaagtgcaaatagtggagcaggagcaggagcaggagcaatattaagtagagctagcATAGACCCTCGAAAAGATTGGTTAAAGAGAATATTGTTGAATAATAACACGAACGATTTAAGGATGTTTTTAAAGGCTAGCGAATTAAAAACAAAAGAAGGTGATTATTGTATGACATGCAAAAGTAACGTAAAACAGTTATTATATCTTCATACAAAAAAGAATTATTGTCATTTATGTGAAGAAATCTTTTGTGCTTATTGTGTAAAAAGTATTGATTTTATGAAAGATGAAAAAGAGAAATATATAAAAATAAGATTATGTAGAAATTGTTTTATATATATAAATGAACTAAAATATATAATTAATCCATAACCCGGGTCGAGGGATATGGCAGCTTAATGTTCGTTTTTCTTATTTATATATTTATACCAATTGATTGTATTTATAACTGTAAAAATGTGTATGTTGTGTGCATATTTTTTTTTGTGCATGCACATGCATGTAAATAGCTAAAATTATGAACATTTTATTTTTTGTTCAGAAAAAAAAAACTTTACACACATAAAATGGCTAGTATGAATAGCCATATTTTATATAAATTAAATCCTATGAATTTATGACCATATTAAAAATTTAGATATTTATGGAACATAATATGTTTGAAACAATAAGACAAAATTATTATTATTATTATTATTTTTACTGTTATAATTATGTGTCTCCTTCAATGATTCATAAATAGTTGGACTTGATTTTTAAAATGTTTATAATATGATTAGCATAGTTAAATAAAAAAAGTTGAAAAATTAAAAAAAAACATATAAACACAAATGATGGTTTTTCCTTCAATTTCGATATCAATTTATAGAAACAAAATATATACTTGTATAATTTTATTTTTTTATATAAATCATTACATATATAATTATACAATATTTTTTCTAAGAGATAATTATATATTAATATATATAAAAAAAGGTGTTTTTTTTTTTTTTTTTTATTTTTATTTTTATTTTATGGTAATATTTTATTTTCCTTATTTTATAAATTATATTAGTTTATATGTGATTAATTTTATATATTATCAATTTATATATTTTTAAATGCTTACTTAATTATCTTTTTTTTTTTTTTTTTTTTTTTTTCCCCTCTTTTTATATTAATTTATTTTTGAAAAAATTGATATATATATATATATATAATATATATATATACATGTAGTAGTATTAAACAATGTATAATATATATAAATAATATATTTATATATTTCATTTCAATTTTAATTTTTTTTGGTTTTTTTTTTTTTTCTTTTTGTCATATTTAAAAAAAATTATATTCATATAAGTTATGCATTTTTTATAAACATTATTCAATATATGTATAATATAATATATATATATATATTAATGTATTATTCCAATGTGCATGATAAAAGAAAAAAATAATATTTATAAAAAAAAAGAAAAATAAAACAAAAAAAGAAAAAAAAAAAAAAAAAAAAAAAAATACAAAAATAAATAATATAATTTATAATTATATATTCTTGTCACAATAAAAATATATATATATATATATATATTTATAATATGTATATTTTAAACTAGAAAAGGAATAACTAATATTTTATTTATTATCATTCAAGATTTATATTTTATAATAATAAATACCTAATAGAAATATATCAGGATCCATGACAGCCAGTTTAACTACCAAGTTCTTGAACAATACCTATGAAAACCCATTTATGAATGCATCCGGTGTTCATTGCATGACTACACAAGAATTAGATGAATTAGCAAACTCTAAAGCTGGCGCATTCATTACAAAGAGTGCTACAACCTTAGAAAGAGAAGGTAACCCTGAACCACGTTACATTTCTGTCCCTCTAGGCAGTATCAACTCCATGGGTTTACCAAACGAAGGTATCGACTACTATTTGTCCTATGTATTAAACCGTCAAAAGAATTATCCTGATGCACCTGCTATTTTCTTCTCAGTTGCTGGTATGAGCATTGATGAAAATTTAAATTTGTTGAGGAAAATCCAAGATAGCGAATTCAACGGTATTACCGAGTTAAACTTGTCTTGTCCTAATGTGCCTGGGAAACCACAAGTTGCTTATGACTTTGACTTGACAAAGGAAACCTTGGAAAAGGTTTTTGCCTTTTTCAAAAAACCTCTTGGTGTCAAGTTGCCTCCTTATTTTGATTTTGCCCATTTTGATATCATGGCAAAAATATTGAACGAGTTCCCATTAGCTTATGTCAACTCTATCAATAGTATAGGAAATGGTCTTTTCATTGATGTGGAGAAGGAGAGTGTAGTAGTGAAGCCAAAGAATGGTTTCGGGGGTATTGGAGGTGAATATGTTAAGCCAACCGCGCTCGCCAATGTTCGTGCATTTTACACTCGTTTGAGACCTGAAATCAAAGTTATCGGTACAGGTGGAATTAAGTCCGGTAAGGATGCATTTGAACATCTTCTATGTGGTGCCTCTATGCTACAGATTGGTACAGAATTACAAAAAGAGGGCGTCAAGATTTTTGAACGTATCGAAAAAGAATTAAAAGACATAATGGAAGCTAAGGGTTATACATCCATAGATCAGTTCCGTGGGAAGTTGAACAGCATTTAAAAGCTTATTTAATAATAGATTAAAAATATTATAAAAATAAAAACATAAACACAGAAATTACAAAAAAAATACATATGAATTTTTTTTTTGTAATCTTCCTTATAAATATAGAATAATGAATCATATAAAACATATCATTATTCATTTATTTACATTTAAAATTATTGTTTCAGTATCTTTAATTTATTATGTATATATAAAAATAACTTACAATTTTATTAATAAACAATATATGTTTATTAATTCATGTTTTGTAATTTATGGGATAGCGATTTTTTTTACTGTCTGTATTTTTCTTTTTTAATTATGTTTTAATTGTATTTTATTTTTATTATTGTTCTTTTTATAGTATTATTTTAAAACAAAATGTATTTTCTAAGAACTTATAATAATAATAAATATAAATTTTAATAAAAATTATATTTATCTTTTACAATATGAACATAAAGTACAACATTAATATATAGCTTTTAATATTTTTATTCCTAATCATGTAAATCTTAAATTTTTCTTTTTAAACATATGTTAAATATTTATTTCTCATTATATATAAGAACATATTTATTAAATCTAGAATTCTATAGTGAGTCGTATTACAATTCACTGGCCGTCGTTTTACAACGTCGTGACTGGGAAAACCCTGGCGTTACCCAACTTAATCGCCTTGCAGCACATCCCCCTTTCGCCAGCTGGCGTAATAGCGAAGAGGCCCGCACCGATCGCCCTTCCCAACAGTTGCGCAGCCTGAATGGCGAATGGCGCCTGATGCGGTATTTTCTCCTTACGCATCTGTGCGGTATTTCACACCGCATATGGTGCACTCTCAGTACAATCTGCTCTGATGCCGCATAGTTAAGCCAGCCCCGACACCCGCCAACACCCGCTGACGCGCCCTGACGGGCTTGTCTGCTCCCGGCATCCGCTTACAGACAAGCTGTGACCGTCTCCGGGAGCTGCATGTGTCAGAGGTTTTCACCGTCATCACCGAAACGCGCGAGACGAAAGGGCCTCGTGATACGCCTATTTTTATAGGTTAATGTCATGATAATAATGGTTTCTTAGACGTCAGGTGGCACTTTTCGGGGAAATGTGCGCGGAACCCCTATTTGTTTATTTTTCTAAATACATTCAAATATGTATCCGCTCATGAGACAATAACCCTGATAAATGCTTCAATAATATTGAAAAAGGAAGAGTATGAGTATTCAACATTTCCGTGTCGCCCTTATTCCCTTTTTTGCGGCATTTTGCCTTCCTGTTTTTGCTCACCCAGAAACGCTGGTGAAAGTAAAAGATGCTGAAGATCAGTTGGGTGCACGAGTGGGTTACATCGAACTGGATCTCAACAGCGGTAAGATCCTTGAGAGTTTTCGCCCCGAAGAACGTTTTCCAATGATGAGCACTTTTAAAGTTCTGCTATGTGGCGCGGTATTATCCCGTATTGACGCCGGGCAAGAGCAACTCGGTCGCCGCATACACTATTCTCAGAATGACTTGGTTGAGTACTCACCAGTCACAGAAAAGCATCTTACGGATGGCATGACAGTAAGAGAATTATGCAGTGCTGCCATAACCATGAGTGATAACACTGCGGCCAACTTACTTCTGACAACGATCGGAGGACCGAAGGAGCTAACCGCTTTTTTGCACAACATGGGGGATCATGTAACTCGCCTTGATCGTTGGGAACCGGAGCTGAATGAAGCCATACCAAACGACGAGCGTGACACCACGATGCCTGTAGCAATGCCAACAACGTTGCGCAAACTATTAACTGGCGAACTACTTACTCTAGCTTCCCGGCAACAATTAATAGACTGGATGGAGGCGGATAAAGTTGCAGGACCACTTCTGCGCTCGGCCCTTCCGGCTGGCTGGTTTATTGCTGATAAATCTGGAGCCGGTGAGCGTGGGTCTCGCGGTATCATTGCAGCACTGGGGCCAGATGGTAAGCCCTCCCGTATCGTAGTTATCTACACGACGGGGAGTCAGGCAACTATGGATGAACGAAATAGACAGATCGCTGAGATAGGTGCCTCACTGATTAAGCATTGGTAACTGTCAGACCAAGTTTACTCATATATACTTTAGATTGATTTAAAACTTCATTTTTAATTTAAAAGGATCTAGGTGAAGATCCTTTTTGATAATCTCATGACCAAAATCCCTTAACGTGAGTTTTCGTTCCACTGAGCGTCAGACCCCGTAGAAAAGATCAAAGGATCTTCTTGAGATCCTTTTTTTCTGCGCGTAATCTGCTGCTTGCAAACAAAAAAACCACCGCTACCAGCGGTGGTTTGTTTGCCGGATCAAGAGCTACCAACTCTTTTTCCGAAGGTAACTGGCTTCAGCAGAGCGCAGATACCAAATACTGTCCTTCTAGTGTAGCCGTAGTTAGGCCACCACTTCAAGAACTCTGTAGCACCGCCTACATACCTCGCTCTGCTAATCCTGTTACCAGTGGCTGCTGCCAGTGGCGATAAGTCGTGTCTTACCGGGTTGGACTCAAGACGATAGTTACCGGATAAGGCGCAGCGGTCGGGCTGAACGGGGGGTTCGTGCACACAGCCCAGCTTGGAGCGAACGACCTACACCGAACTGAGATACCTACAGCGTGAGCTATGAGAAAGCGCCACGCTTCCCGAAGGGAGAAAGGCGGACAGGTATCCGGTAAGCGGCAGGGTCGGAACAGGAGAGCGCACGAGGGAGCTTCCAGGGGGAAACGCCTGGTATCTTTATAGTCCTGTCGGGTTTCGCCACCTCTGACTTGAGCGTCGATTTTTGTGATGCTCGTCAGGGGGGCGGAGCCTATCGAAAAACGCCAGCAACGCGGCCTTTTTACGGTTCCTGGCCTTTTGCTGGCCTTTTGCTCACATGTTCTTTCCTGCGTTATCCCCTGATTCTGTGGATAACCGTATTACCGCCTTTGAGTGAGCTGATACCGCTCGCCGCAGCCGAACGACCGAGCGCAGCGAGTCAGTGAGCGAGGAAGCGGAAGAGCGCCCAATACGCAAACCGCCTCTCCCCGCGCGTTGGCCGATTCATTAATGCAGCTGGCACGACAGGTTTCCCGACTGGAAAGCGGGCAGTGAGCGCAACGCAATTAATGTGAGTTAGCTCACTCATTAGGCACCCCAGGCTTTACACTTTATGCTTCCGGCTCGTATGTTGTGTGGAATTGTGAGCGGATAACAATTTCACACAGGAAACAGCTATGACCATGATTACGCCAAGCTATTTAGGTGACACTATAGAATACTC
